# Modelling the distribution of rare invertebrates by correcting class imbalance and spatial bias

**DOI:** 10.1101/2022.01.03.474856

**Authors:** Willson Gaul, Dinara Sadykova, Hannah J. White, Lupe León-Sánchez, Paul Caplat, Mark C. Emmerson, Jon M. Yearsley

## Abstract

**Aim:** Soil arthropods are important decomposers and nutrient cyclers, but are poorly represented on national and international conservation Red Lists. Opportunistic biological records for soil invertebrates are often sparse, and contain few observations of rare species but a relatively large number of non-detection observations (a problem known as class imbalance). Robinson et al. (2018) proposed a method for sub-sampling non-detection data using a spatial grid to improve class balance and spatial bias in bird data. For taxa that are less intensively sampled, datasets are smaller, which poses a challenge because under-sampling data removes information. We tested whether spatial under-sampling improved prediction performance of species distribution models for millipedes, for which large datasets are not available. We also tested whether using environmental predictor variables provided additional information beyond what is captured by spatial position for predicting species distributions.

**Location:** Island of Ireland.

**Methods:** We tested the spatial under-sampling method of Robinson et al. (2018) by using biological records to train species distribution models of rare millipedes.

**Results:** Using spatially under-sampled training data improved species distribution model sensitivity (true positive rate) but decreased model specificity (true negative rate). The decrease in specificity was minimal for rarer species and was accompanied by substantial increases in sensitivity. For common species, specificity decreased more, and sensitivity increased less, making spatial under-sampling most useful for rare species. Geographic coordinates were as good as or better than environmental variables for predicting distributions of two out of six species.

**Main Conclusions:** Spatial under-sampling improved prediction performance of species distribution models for rare soil arthropod species. Spatial under-sampling was most effective for rarer species. The good prediction performance of models using geographic coordinates is promising for modeling distributions of poorly studied species for which little is known about ecological or physiological determinants of occurrence.

## 1 Introduction

Biological records datasets contain relatively few records of rare species, because rare species generally have lower abundances and occur in fewer locations than common species. When rare species have behavioral or physical characteristics (e.g. small size) that make them difficult to find or identify, they will be even less well represented in datasets. Two common problems for modelling rare species distributions using biological records are class imbalance (He & Garcia, 2009) and spatial bias in the data. Spatial bias is pervasive at many spatial scales in biological records data for all taxa (Amano & Sutherland, 2013; Oliveira et al., 2016). Class imbalance – usually in the form of a preponderance of non-detection observations and few detections of the focal species – is a problem more restricted to species that are rare, difficult to find, or difficult to identify.

Robinson et al. (2018) proposed a method of spatially under-sampling opportunistic species occurrence data in a way that improves both class balance and spatial bias in data before modelling distributions of rare bird species. Their innovation was to use a spatial grid to filter only non-detection data, of which there is plenty, while keeping all detection data, of which there is little for rare species. This approach improves both class balance and spatial bias in the data, and improves upon previous attempts to filter or sub-sample data to address spatial bias, which risk removing too much information about rare species. Robinson et al. (2018) demonstrated spatial under-sampling by modelling the distribution of a rare bird species in California, USA, using eBird data (https://ebird.org). We applied their method to model distributions of six millipede species in Ireland using a dataset an order of magnitude smaller. Few invertebrates (with the possible exception of butterflies) will ever have datasets as large as those available for birds. To the best of our knowledge, ours is the first test of spatial under-sampling to improve species distribution model (SDM) predictions using such a small dataset, and thus provides important insight about how relevant this method is for non-charismatic, poorly recorded taxa.

Invertebrates are poorly represented in conservation research (Donaldson et al., 2016) and on national and international lists of threatened and endangered species, including the IUCN Red List (Cardoso et al., 2011; Cardoso et al., 2012). Invertebrates play key roles in many ecosystem processes, including decomposition and nutrient cycling in soil (Bardgett, 2005; Bardgett & Wardle, 2010), pollination (Potts et al., 2016), and structuring ecosystems (Risch et al., 2018). Evaluations of extinction risk depend in part on knowledge of species distributions (IUCN, 2012). Species distribution modelling methods that produce reliable predicted distributions can aid conservation threat assessment (Cardoso et al., 2011; Maes et al., 2015). For invertebrates, this will often require modelling distributions using data sets much smaller than the datasets available for more easily recorded taxa such as birds.

Biological records data for millipedes in Ireland are not nearly as extensive as data for some other taxa such as birds and vascular plants, but there have been two relatively intense periods of millipede recording in Ireland, culminating in a millipede distribution atlas (Lee, 2006). The data were vetted (and largely collected) by regional experts (Lee, 2006), so the dataset is of high taxonomic quality.

For millipedes in our study area, the direct environmental drivers of species occurrence (Austin, 2007) are not well known. Lee (2006) provided a comprehensive analysis of habitat affinities for millipedes in Great Britain and Ireland based on habitat data recorded as part of a British and Irish millipede recording scheme. But sampling in the recording scheme was opportunistic, and low numbers of records for some habitats made inference about habitat associations imprecise (Lee, 2006). The associations examined in Lee (2006) were largely land use, habitat, and soil characteristics - climatic variables were not tested. Kime (1999, 2001, 2004) discussed the environmental determinants of millipede species distributions in the UK, Ireland, and continental Europe using descriptive evaluations of locations of records and distribution patterns, but did not perform statistical analyses. Previous analyses (Kime, 1999; Kime, 2001; Kime, 2004; Lee, 2006) therefore provide only a starting point for identifying the direct drivers of Irish millipede species distributions; it seems likely that those studies did not identify all important environmental drivers, and it is possible that some drivers were incorrectly identified as important.

Even with well-studied taxa, the ability of modelers and models to identify biologically meaningful environmental predictors is limited (Beale, Lennon & Gimona, 2008; Currie, Pétrin & Véronique, 2020). In a study of North American breeding birds, Bahn and McGill (2007) found that SDMs using only geographic coordinates performed better than models using environmental predictors. Fourcad et al. (2017) found that distributions of European Red Listed species were predicted nearly as well when the color values of paintings were used as predictors as when environmental predictors were used. This suggests that environmental variables might not be capturing anything more than spatial autocorrelation from sources that are either exogenous (e.g. correlation in environmental drivers) or endogenous (e.g. dispersal) to the modeled species. The ability to accurately predict species distributions using only spatial information (Bahn & McGill, 2007) presents an opportunity for predictive modelling of taxa such as millipedes, for which physiological and ecological knowledge is poor. In contrast, the reliability of inferences about the effects of environmental drivers of species distributions are cast into doubt if models using purely spatial information predict distributions as well as models using environmental variables.

We modeled the distribution of six millipede species in Ireland using biological records and the spatial under-sampling method of Robinson et al. (2018). We asked the following questions. 1) Does spatially under-sampling training data improve the predictive performance of SDMs for rare millipedes? 2) Does the effectiveness of using spatially under-sampled training data depend on the rarity of the species? 3) Do models using environmental predictors have better prediction performance than models using geographic coordinates as predictor variables? 4) Does spatially under-sampling training data change the apparent relative importance of predictor variables compared to models trained with raw data?

## 2 Methods

We modeled the distribution of rare millipedes in Ireland using random forests (Breiman, 2001), which can model non-linear relationships and interactions between predictor variables, and were used by Robinson et al. (2018). To determine what types of information (environmental, spatial, or seasonal) were important for predicting millipede detections, we trained four types of models: 1) a seasonal base model (*SEASON + LIST LENGTH*) that predicted millipede occurrence records as a function of month and a “checklist length” variable (details below) that we expected would capture variability in sampling effort among checklists; 2) a spatial model (*COORDINATES + SEASON + LIST LENGTH*) that used the base model plus geographic coordinates (eastings and northings of the TM75 Irish Grid Reference system); 3) an environmental model (*ENVIRONMENT + SEASON + LIST LENGTH*) that used the base model plus environmental covariates; and 4) an environmental and spatial model (*ENVIRONMENT + COORDINATES + SEASON + LIST LENGTH*) that used the base model plus environmental covariates and geographic coordinates.

We trained all models with both raw data, in which there was considerable class imbalance and spatial bias, and with spatially under-sampled data (Robinson et al., 2018) in which the class imbalance and spatial bias were partially corrected (Fig. 1, Figs. S1-S2).

**Figure 1:**
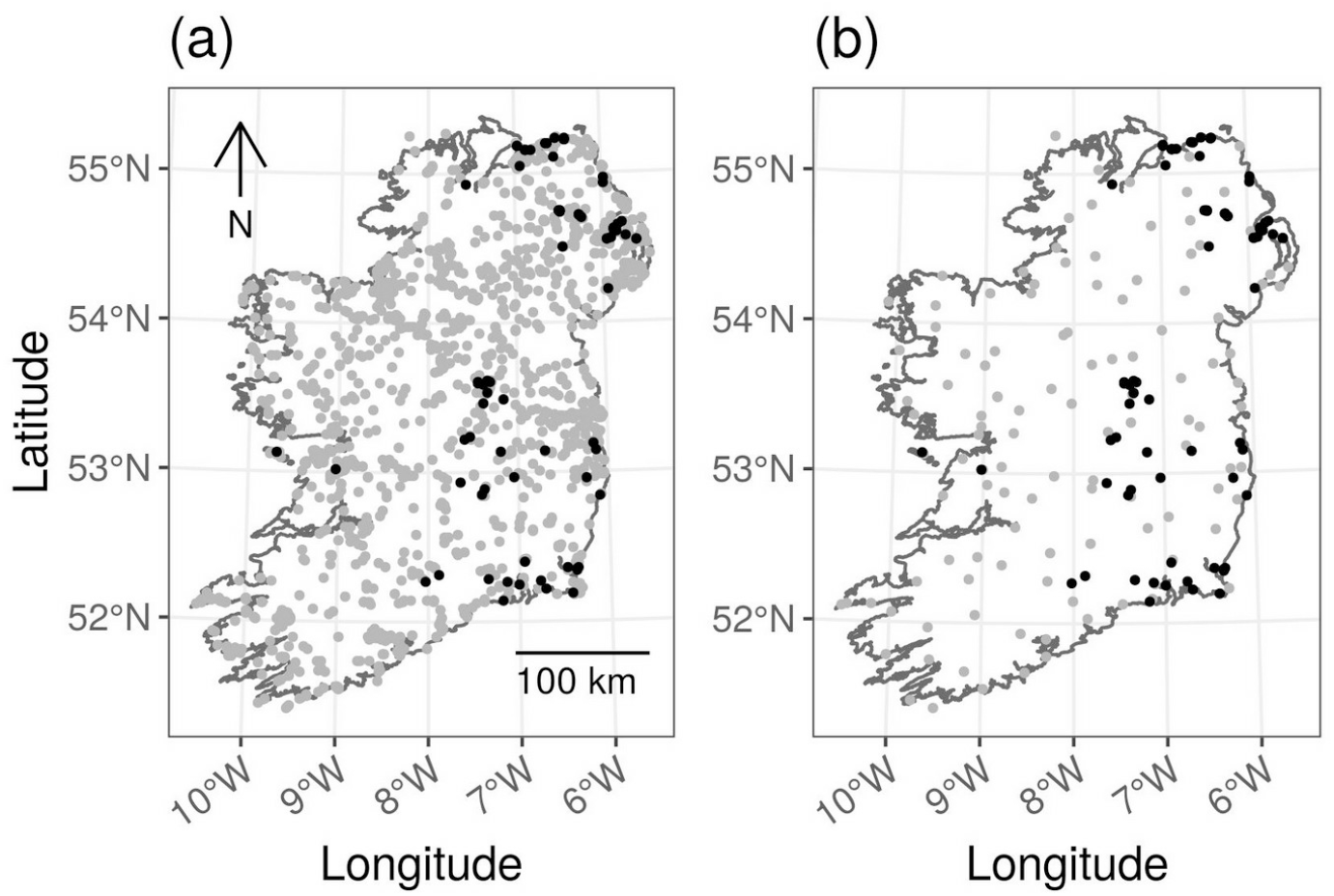
Raw (a) and spatially under-sampled (b) observation data for the millipede *Ommatoiulus sabulosus* on the island of Ireland. Spatial under-sampling involved keeping all checklists on which the species was detected (black points), but spatially filtering the non-detection checklists (grey points) by randomly choosing only a single non-detection checklist from each cell of a 30 × 30 km grid that was randomly positioned over the study extent (grid not shown on these figures). Spatial under-sampling improved class balance and reduced the spatial bias of the non-detection data.

### 2.1 Study Species

We modeled distributions of four rare millipede species detected on fewer than 10% of checklists: *Macrosternodesmus palicola* Brölemann, 1908; *Boreoiulus tenuis* (Bigler, 1913); *Ommatoiulus sabulosus* (Linnaeus, 1758); and *Blaniulus guttulatus* (Fabricius, 1798), and two more common species: *Glomeris marginata* (Villers, 1789) and *Cylindroiulus punctatus* (Leach, 1815). *C. punctatus* was the most commonly recorded species in our dataset (Table 1). Three of the species (*M. palicola, Boreoiulus tenuis*, and *Blaniulus guttulatus*) are believed to be somewhat synanthropic, while the other three *O. sabulosus, G. marginata*, and *C. punctatus*) are not (Lee, 2006).

**Table 1:** Environmental predictor variables used to model millipede species in Ireland. For each millipede species, the number of positive detections with a spatial precision of 1 km or less is shown, along with the proportion of all checklists on which the species was detected. The environmental predictor variables used for each species are indicated with an “X”.

| Species | Number of detections | Proportion of checklists with a detection | mean annual minimum temperature | mean annual precipitation | elevation | artificial surfaces land cover | arable land cover | wetlands land cover | forest and semi-natural areas land cover | pasture land cover |
| --- | --- | --- | --- | --- | --- | --- | --- | --- | --- | --- |
| *Macrosternodesmus palicola* | 61 | 0.035 |  | X |  | X | X |  |  |  |
| *Boreoiulus tenuis* | 73 | 0.042 |  | X | X | X | X |  |  |  |
| *Ommatoiulus sabulosus* | 77 | 0.044 |  | X | X | X |  | X |  |  |
| *Blaniulus guttulatus* | 143 | 0.081 | X | X | X | X | X |  | X |  |
| *Glomeris marginata* | 325 | 0.185 | X | X | X | X | X |  | X |  |
| *Cylindroiulus punctatus* | 615 | 0.350 | X | X | X | X | X |  | X | X |

All of the species we modeled take multiple years to reach maturity in our study area, so they should be present year round. However, millipedes in Ireland show seasonal patterns of activity and detectability. For example, many species move deeper into leaf litter or soil to avoid cold temperatures or dry conditions (Lee, 2006). Differences in maturation speeds, life spans, and activity patterns between species (and between sexes within species) mean that the number of individuals, number of adults, and relative proportions of the sexes may not be constant at all times of year. Adults are generally easier to identify than juveniles. For some species only sexually mature males can be identified to species level based on morphology (i.e. without molecular evidence), which means both the total number of species that can be identified and recorded, and the probability of recording any particular species, are likely to change over the course of a year in any given location.

### 2.2 Millipede occurrence data

We downloaded records of all millipedes (including but not limited to our six focal species) on the island of Ireland for the years 1971 to 2020 from the Global Biodiversity Information Facility (GBIF.org, 2021). The data included records from multiple sources, including the British Myriapod and Isopod Group recording scheme (Biological Records Centre, 2017; Lee, 2006). The data were presence-only records of millipede species occurrence, and did not include explicit sampling effort, sampling method, or non-detection information. We grouped records into recording event “checklists”, where a checklist was defined as a unique combination of date, location, and observer, and each species was either detected or not detected (van Strien et al., 2010). We calculated checklist length by counting the number of species detected on each checklist. We retained for analysis only checklists with spatial precision of 1 km or less (n = 1757 checklists).

### 2.3 Environmental data

The species we selected are believed to respond to a variety of land cover and habitat characteristics, soil types, and human disturbance (Lee, 2006). We identified (when possible) remotely sensed environmental variables that corresponded to the strongest habitat affinities reported for each focal species in Lee (2006). All six of our focal species were reported to respond strongly to urban vs. rural land use classifications in Lee (2006). Four species showed strong relationships with woodland land cover in Lee (2006). At least one species, *G. marginata*, may be unable to tolerate low temperatures (Lee, 2006), which might exclude it from higher elevations in Ireland.

The environmental variables we used included elevation, two climate variables, and five land cover variables (Table 1). We calculated the value of each predictor variable in 1 × 1 km grid cells covering Ireland. We used the mean elevation of each grid square (calculated by interpolating using ordinary kriging) from the ETOPO1 Global Relief Model (Amante & Eakins, 2009). For the land cover variables, we calculated the proportion of each grid cell covered by “artificial surfaces”, “forest and semi-natural areas”, “wetlands”, “pasture”, and “arable land” classes from the CORINE Land Cover database (CORINE, 2012). We downloaded gridded climate variables from the E-OBS European Climate Assessment and Dataset EU project (Haylock et al., 2008; van den Besselaar et al., 2011), and, for each 1 km^2^ grid cell, calculated the mean annual precipitation for the years 1995 to 2016 (excluding years 2010 through 2012 because of missing data), and the mean annual low temperature across years 1995 to 2016. (Annual low temperature was taken to be the 2% quantile of daily temperatures from a year. The 2% quantile was used to prevent erroneous extreme data values from influencing the results). Low temperature was used rather than other temperature variables because Kime (1999) expected low temperature to be an important determinant of millipede distribution in northern Europe, while high summer temperature was expected to be an important determinant in southern Europe. Cold winters are believed to limit the distribution of *G. marginata* (Kime, 2004), though other species have behavioral responses (e.g. burrowing in soil or dead wood) that allow them to survive cold periods (Kime, 2004).

### 2.4 Spatial under-sampling

Our spatial under-sampling process followed Robinson et al. (2018). We first split the millipede checklists into two groups: non-detection checklists, on which the focal species was not detected, and detection checklists, on which the focal species was detected. We then generated a randomly positioned 30 km x 30 km grid over the study extent using the ‘blockCV’ R package (Valavi et al., 2019). We sub-sampled the non-detection checklists by randomly selecting one non-detection checklist from each 30 km x 30 km grid cell (provided a grid cell contained at least one non-detection checklist). We kept all detection checklists. We then combined the spatially under-sampled non-detection checklists with the detection checklists to create a spatially under-sampled dataset. We used this spatially under-sampled dataset as training data for SDMs. We evaluated spatial evenness in the raw and spatially under-sampled data by measuring Simpson’s evenness for the number of checklists in 30 km x 30 km grid squares.

All spatial data processing used the ‘sf’, ‘sp’, ‘raster’, ‘rgdal’, ‘gstat’, and ‘tidyverse’ packages in R version 3.6 (Bivand et al., 2018; Gräler et al., 2016; Hijmans, 2018; Pebesma, 2018; R Core Team, 2020; Ross, 2018; Wickham, 2017).

### 2.5 Species distribution models

We trained random forest SDMs with the ‘randomForest’ function in R (Liaw & Wiener, 2002) using three-fold cross-validation (three folds were used because using five folds often resulted in folds with no or few detections, which made model testing difficult or impossible in those folds). The units of analysis were sampling event checklists (n = 1757), on which each species was either detected or not detected (van Strien et al., 2010). We modeled species detections as a function of predictor variables expected to influence species occupancy and/or detectability. We did not separately estimate occupancy and detectability, as is done in hierarchical occupancy/detection models (MacKenzie et al., 2002) because our data contained few repeat survey visits within a year, but we included predictor variables that we expected would primarily influence either occupancy or detectability. Occupancy covariates were geographic coordinates and environmental covariates. The month of each observation was included primarily as a detectability covariate because most millipede species in our study have seasonal changes in behavior and/or seasonal life cycles that make them easier to detect and identify at some times of year. To allow for the periodic variation in detectability with month, we represented months as integers (1 through 12) and used cosine and sine transformations of month to create two separate transformed month variables that we provided to the random forest SDMs (James, 2011; see Appendix S1). Checklist length was used as a proxy for sampling effort (Szabo et al., 2010; Isaac et al., 2014) and was thus primarily a covariate for detectability, though checklist length likely also varied with environmental conditions and species richness in our study (see Discussion), and is therefore potentially related to occupancy as well as detectability. We expected the probability of detecting each focal species to increase with checklist length.

We used different environmental predictor variables for each species based on the environmental and habitat affinities reported in Lee (2006). The limited amount of occurrence data available for rare millipede species made over-fitting a concern. Each model included variables indicating the month in which the record was collected, and checklist length. For models including environmental covariates, we selected the number of predictor variables to use in each model based on the number of positive detections for each species, so that there were at least ten detections of the focal species per predictor variable (Table 1).

Each checklist was randomly assigned to one of three cross-validation folds. We used random forests for classification to predict the detection or non-detection of the focal species on each checklist. For each random forest model, we grew 1000 trees with a terminal node size of one, using the largest integer less than the square root of the number of predictor variables as the number of variables to consider for splitting at each split. For each species, we performed 33 modelling runs, each time fitting models with three-fold cross-validation. This produced a total of 99 fitted models for each species with each combination of training data type (raw or spatially under-sampled) and model type (*SEASON + LIST LENGTH, COORDINATES + SEASON + LIST LENGTH, ENVIRONMENT + SEASON + LIST LENGTH*, and *ENVIRONMENT + COORDINATES + SEASON + LIST LENGTH*). We generated a new spatially under-sampled dataset for each of the 33 model fitting iterations.

We assessed prediction performance of each fitted model by predicting to checklists in the cross-validation test fold. We measured the ability of models to accurately discriminate between detections and non-detections using the area under the receiver operating characteristic curve (AUC) (Fielding & Bell, 1997), Cohen’s Kappa (Cohen, 1960), and sensitivity (true positive rate) and specificity (true negative rate) at the threshold that maximized Cohen’s Kappa. Sensitivity measured the ability of models to correctly predict which checklists had detections, while specificity measured the ability of models to correctly predict which checklists had non-detections. We also measured the calibration of the predicted probabilities of models using the Brier score (Brier, 1950) following Robinson et al. (2018). We calculated prediction performance measures on spatially under-sampled test datasets, because the goal of SDMs was to predict species occurrence at all locations in Ireland, where all locations are equally important (Robinson et al., 2018). Spatial under-sampling test data reduces the spatial bias in the data, and therefore reduces the extent to which prediction performance measures are dominated by how models predict in the most heavily sampled areas (Fink et al., 2010; Robinson et al., 2018).

We averaged the variable importance (mean decrease in node impurity over all trees due to splitting on each variable, measured using the Gini index) and partial dependence measures produced by random forest models over all 99 iterations of each model (see Appendix S1). We made predicted distribution maps for each species using the most complex model (*ENVIRONMENT + COORDINATES + SEASON + LIST LENGTH*), fixing the checklist length at two (the median in the observed data) and generating predictions for every month. We then averaged the monthly predictions over the entire annual cycle from all 99 model iterations to get the average predicted probability of detecting the focal species on a checklist of length two for each grid cell. The variability in model predictions for each grid cell was visualized by mapping the standard error of the mean annual predictions from the 99 model iterations (Fink et al., 2014). The standard errors show the variability in the mean prediction due to differences in which data were included the cross-validation training sets, but do not show variation in predictions due to changes over the annual cycle.

## 3 Results

### 3.1 Spatial under-sampling

Spatial under-sampling improved both the class balance (Fig. S1) and spatial evenness (Fig. S2) of training data for all species. Using spatially under-sampled training data generally improved overall discrimination performance (AUC) for rarer species (Fig. 2), though there was minimal or no improvement for some models of the rarest species (*M. palicola*, Fig. 2). For the two more common species (*G. marginata* and *C. punctatus*), spatial under-sampling did not improve prediction performance as much as it did for rarer species, and notably reduced the overall performance of the simplest model (Fig. 2).

**Figure 2:**
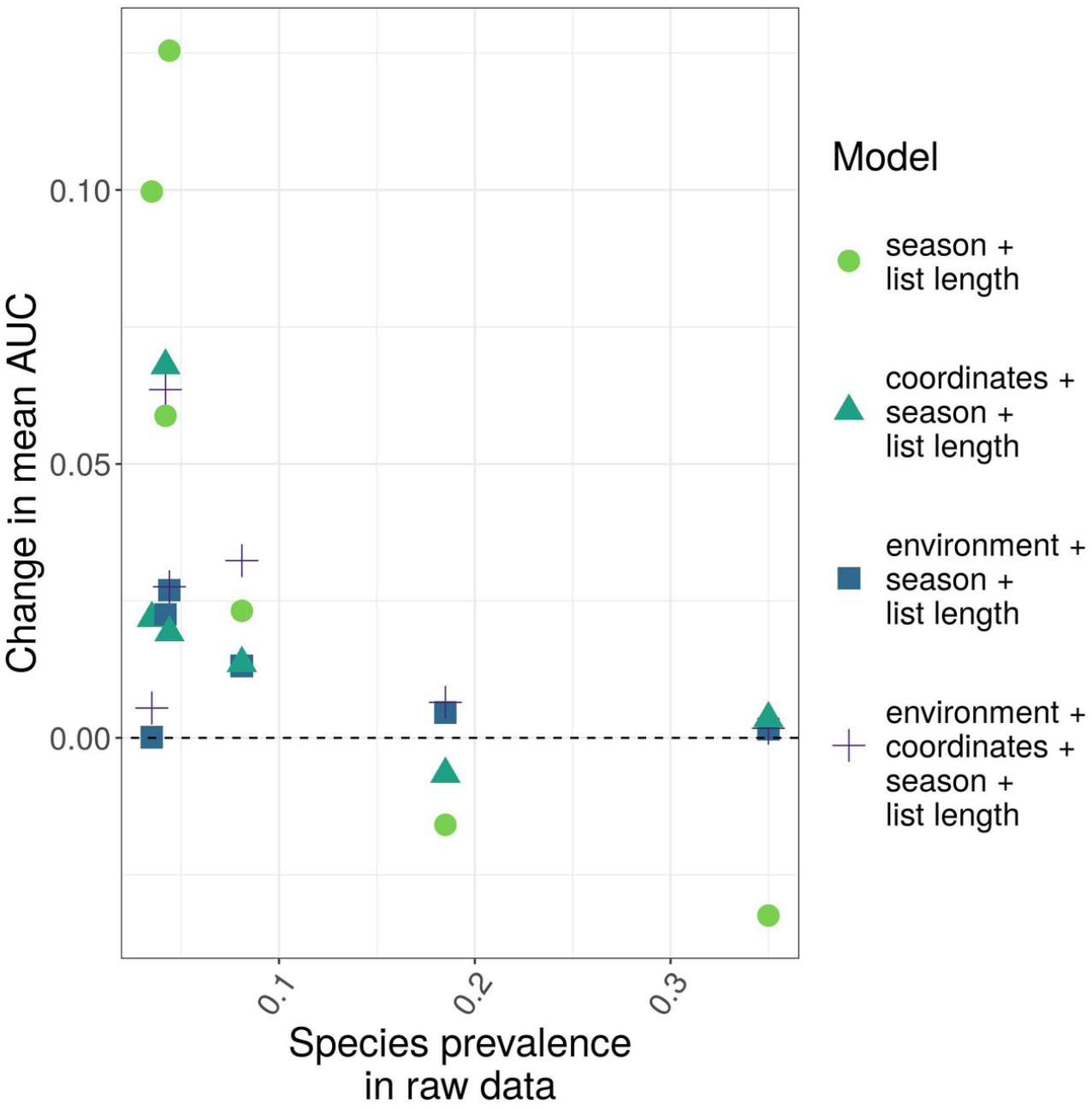
Change in mean prediction performance as a function of species prevalence in the original data, when species distribution models (SDMs) were trained using spatially under-sampled rather than raw data. Results are shown for random forest SDMs for six millipede species in Ireland. Points above the horizontal dotted line indicate that spatially under-sampling the training data improved model prediction performance. Prediction performance was measured using the area under the receiver operating characteristic curve (AUC). Four models with different sets of predictor variables were tested.

Prediction performance of the most complex model (*ENVIRONMENT + COORDINATES + SEASON + LIST LENGTH*), which was the best model for four species and the second best model for two species (Fig. 3), was generally improved by spatially under-sampling training data according to most performance metrics, including threshold-dependent (Kappa, sensitivity) and -independent (AUC) discrimination metrics, and Brier score (Fig. 4). Sensitivity was notably improved by spatially under-sampling training data. Spatial under-sampling reduced model specificity. The decrease in specificity with spatially under-sampled training data was greatest for the most common species and smallest for the rarer species (Fig. 4). Thus, for rare species, spatial under-sampling seems to be an effective way of increasing model sensitivity without sacrificing too much specificity.

**Figure 3:**
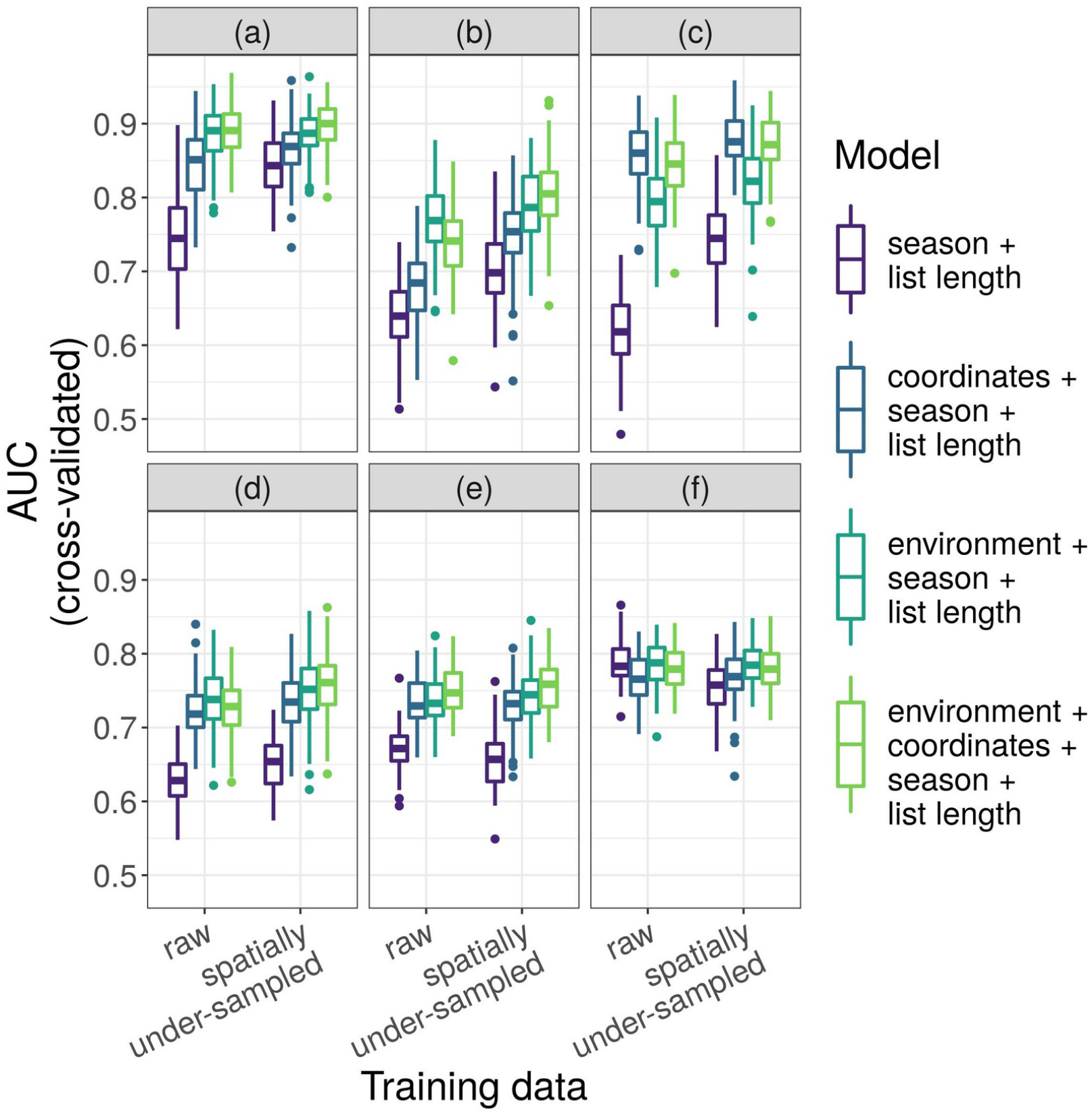
Prediction performance (AUC) of random forest species distribution models for six millipede species in Ireland. Results are shown for models trained with raw and with spatially under-sampled data (left and right box plots, respectively, within each panel). The six modeled species, arranged from rarest to most common in our data, were *Macrosternodesmus palicola* (a), *Boreoiulus tenuis* (b), *Ommatoiulus sabulosus* (c), *Blaniulus guttulatus* (d), *Glomeris marginata* (e), and *Cylindroiulus punctatus* (f). Box plots show the distribution of AUC values for 99 replicates of each model; boxes contain the middle 50% of the data, the horizontal line within each box shows the median.

**Figure 4:**
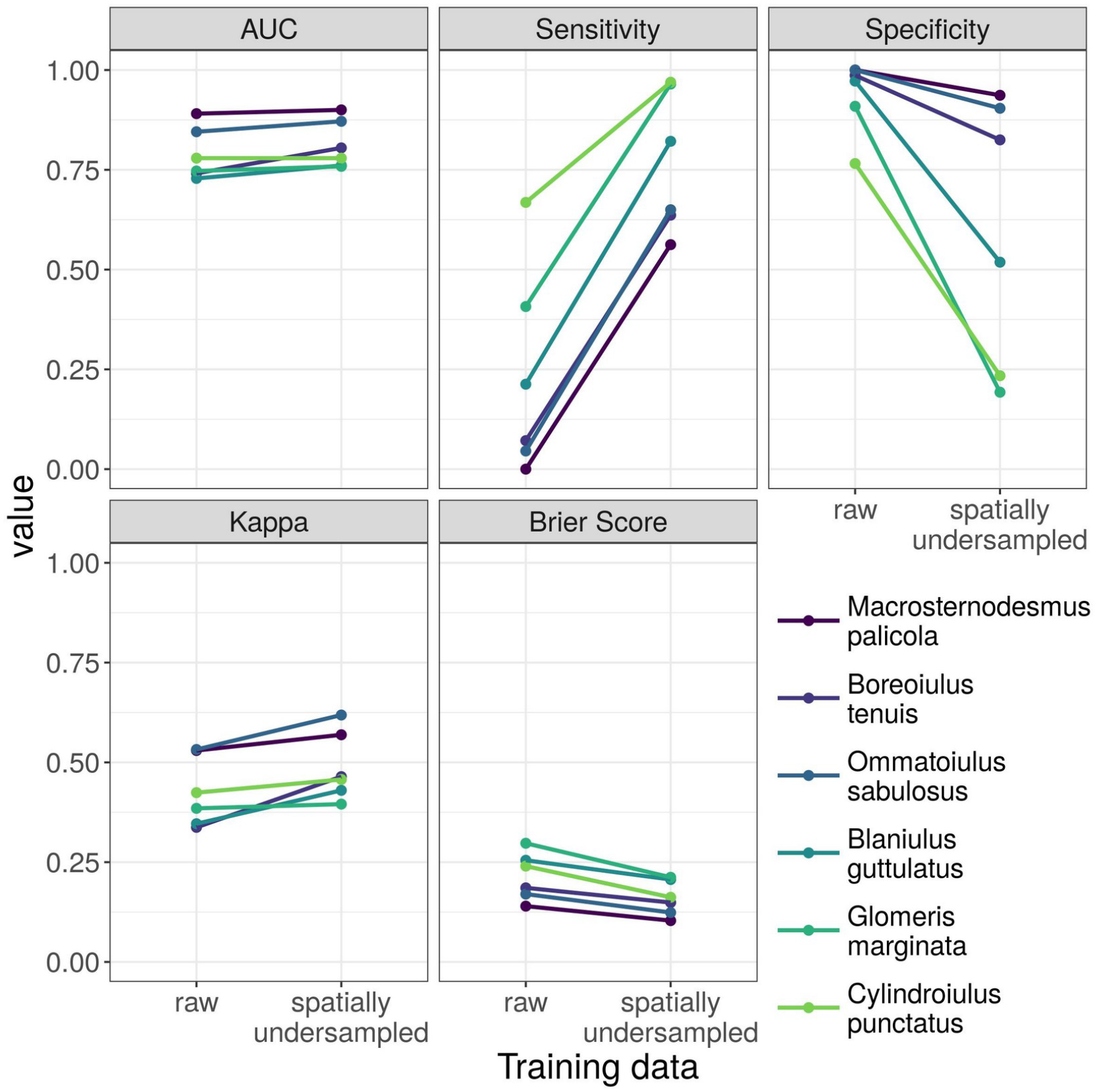
Prediction performance of species distribution models trained with raw (left) and with spatially under-sampled (right) data for six species of millipede in Ireland. Points show the median value of each prediction performance measure from 99 replicate models (33 replicates of three-fold cross-validation) fit to each species with each type of training data. Darker colored points and lines indicate rarer species, and lighter colors indicate more common species. Sensitivity was substantially improved for all species when training data were spatially under-sampled. Spatial under-sampling caused an undesirable decrease in specificity for all species, but the decrease was smaller for rarer species and greater for more common species.

Rankings of variable importance changed when models were trained with spatially under-sampled rather than raw data (Figs. S3-S4). For all species, the least important variable remained the least important regardless of the training data used, but for four out of six species, the most important variable was different depending on whether training data had been spatially under-sampled (Figs. S3-S4).

### 3.2 Environmental vs. spatial models

The spatial model (*COORDINATES + SEASON + LIST LENGTH*) was better than the environmental model (*ENVIRONMENT + SEASON + LIST LENGTH*) for *O. sabulosus* (Fig. 3c), similar to the environmental model for *G. marginata* (Fig. 3e), and worse than the environmental model for the remaining four species (Fig. 3).

The simplest model (*SEASON + LIST LENGTH*) generally performed worse than more complex models that included geographic coordinates, environmental variables, or both (Fig. 3, mean difference in AUC between the simplest and most complex models for each species when using spatially under-sampled data = −0.09, range = −0.13 to −0.03). A notable exception was for the common species *C. punctatus*, for which the *SEASON + LIST LENGTH* model trained with raw data was among the best models (Fig. 3f). This suggests that information about the location of a checklist was not important for predicting the probability of recording *C. punctatus*.

### 3.3 Effects of covariates on species occurrence and detection

Partial dependence plots of the marginal effect of each variable from the most complex model showed plausible relationships. The probability of detecting the focal species on a checklist generally increased in a decelerating curve with checklist length, as expected (Fig. 5f, Figs. S5-S10). Checklist length was among the most important variables for all species except *O. sabulosus*, when assessing variable importance for the most complex model trained with spatially under-sampled data (Figs. S3-S4).

**Figure 5:**
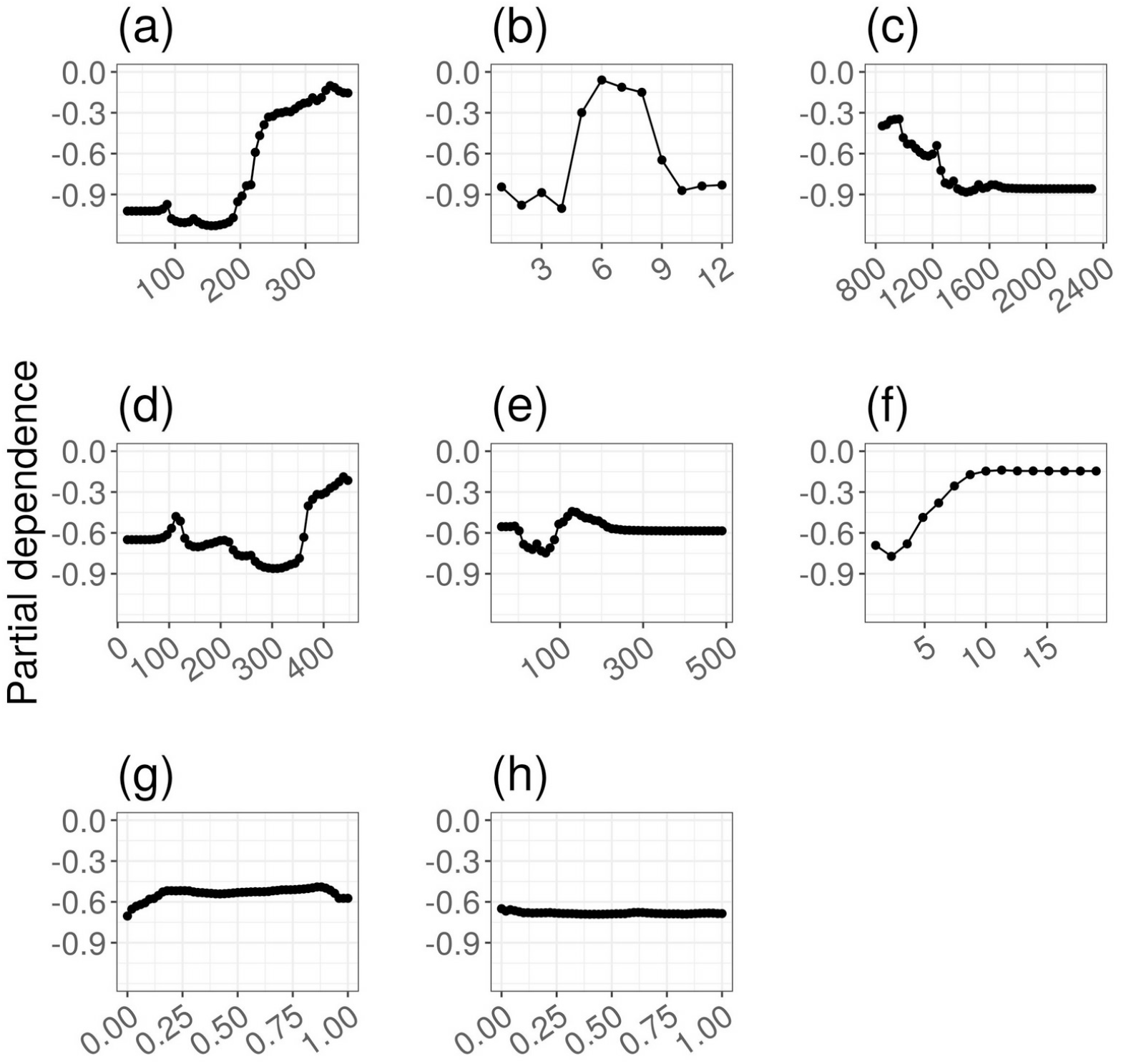
Partial dependence plots showing the effect of predictor variables on the probability of detecting the millipede *Ommatoiulus sabulosus* in Ireland. Results are shown for the *ENVIRONMENT + COORDINATES + SEASON + LIST LENGTH* model trained with spatially under-sampled data. The vertical axes show the partial dependence measure (Supplementary Article S1), with higher values indicating a higher probability of detecting the species. The horizontal axes show the value of the predictor variables: (a) kilometers east of the origin point of the TM75 Irish Grid Reference system; (b) month (one indicates January); (c) annual precipitation (mm); (d) kilometers north of the origin point of the TM75 Irish Grid Reference system; (e) elevation (m); (f) checklist length (number of records); (g) proportion of grid cell area covered by artificial surfaces; and (h) proportion of grid cell area covered by wetlands.

Seasonal changes in the probability of detection were clearly visible in partial dependence plots for the month variable for the three rarest species, with increased probability of detection in winter for *M. palicola* (Fig. S5) and *Boreoiulus tenuis* (Fig. S6), and increased probability of detection in summer for *O. sabulosus* (Fig. 5b, Fig. S7). For *Blaniulus guttulatus*, seasonal patterns of detectability were less clear, but detectability appeared lowest in summer and highest in spring and fall (Fig. S8). There were no clear seasonal patterns in detectability for *G. marginata* (Fig. S9), or *C. punctatus* (Fig. S10).

## 4 Discussion

We set out to answer four questions: 1) Does spatially under-sampling training data improve the predictive performance of SDMs for rare millipedes? 2) Does the effectiveness of spatial under-sampling depend on the rarity of the species? 3) Do models using environmental predictors have better prediction performance than models using geographic coordinates as predictor variables? 4) Does spatially under-sampling training data change the apparent relative importance of predictor variables compared to models trained with raw data? Briefly, the answers to these questions were: 1) yes; 2) yes; 3) usually, but not always; and 4) usually, but not always.

Under-sampling data to address class imbalance has been explored in the machine learning literature, and is used in a wide variety of applied settings (reviewed in Haixiang et al., 2017). The innovation of Robinson et al. (2018) was to perform the under-sampling using a spatial grid to simultaneously improve the spatial evenness of data. We further tested the usefulness of Robinson et al.’s (2018) method for modeling distributions of rare invertebrates using a small dataset. We found that spatial under-sampling can be usefully applied to small datasets. It improved most measures of prediction performance of our random forest SDMs, and was most effective for rarer species.

Spatial under-sampling increased sensitivity at the expense of specificity in our models. Spatial under-sampling will therefore be most useful in applications where false positive predictions are not particularly problematic. High-sensitivity SDMs can guide targeted surveys for rare millipedes, which would help fill knowledge gaps and could facilitate production of a national Red List for Irish millipedes. High sensitivity SDMs could also be used to produce a “short list” of locations that are candidates for conservation or management of focal species. Locations on a “short list” could subsequently be surveyed more intensively to confirm focal species presence. Any false positive predictions (locations in which further surveying revealed that the species is absent despite the model’s positive prediction), could then be excluded, resulting in a final list of locations for conservation action. False positive predictions from models trained with spatially under-sampled data could be problematic if conservation or policy decisions ignore false positives. For example, deciding not to list a species as threatened because a model predicted a large distribution would be a bad decision if that model had high sensitivity and low specificity.

The millipede dataset we used had about 0.02 checklists per km^2^ – an order of magnitude less data than the dataset used by Robinson et al. (2018), which had about 0.26 to 0.71 checklists per km^2^ (for their winter and summer models, respectively). Robinson et al. (2018) reported no loss of specificity when using spatially under-sampled training data. In contrast, models for all of our species had some loss of specificity when using spatially under-sampled training data. Perhaps the limited amount of information available in our smaller dataset meant that there was less “spare” non-detection data to be discarded before model specificity declined. Large reductions in specificity indicated that models trained with spatially under-sampled data over-estimated distributions of the more common species in our study. For rarer species, small losses of specificity and large gains in sensitivity when training data were spatially under-sampled suggested that improving class balance at the cost of discarding non-detection data is worth exploring with small datasets such as ours.

Given the good performance of spatial under-sampling here and in Robinson et al. (2018), it is worth exploring how to optimally tune the procedure. The size of the spatial grid used for under-sampling, and the class balance of the under-sampled dataset could be systematically explored and tuned using cross-validation, as is done for other model parameters in machine learning settings (Hastie et al., 2009). Finer spatial under-sampling grids will result in more non-detection checklists in under-sampled data. If the number of grid cells in the spatial grid is very large compared to the number of presence records, the number of non-detection points will still greatly exceed the number of presence points in the spatially under-sampled data, and class balance will remain poor. We suggest that future studies optimize the spatial grid resolution and class balance by systematically varying the grid resolution, and, optionally, the number of non-detection observations drawn from each grid cell, until out-of-sample predictive performance of SDMs is maximized, or until the number of non-detection observations is small enough that class balance meets some target threshold (e.g. at least 25% of data are detections, as in Robinson et al., 2020).

The spatial model (*COORDINATES + SEASON + LIST LENGTH*) clearly outperformed the environmental model (*ENVIRONMENT + SEASON + LIST LENGTH*) for *O. sabulosus*. For that species, the environmental covariates did not have any more predictive power than did information about spatial location. The distribution of *O. sabulosus* in Ireland might be determined primarily by non-environmental factors, such as dispersal or biotic interactions. Alternatively, it could be that *O. sabulosus* distribution is determined by environmental variables that were not in our model, but which were spatially auto-correlated so that geographic coordinates were effective proxies.

For all species, the model including both environmental variables and geographic coordinates as predictors was among the best models. Because our model evaluations used cross-validation, this is unlikely to be an artifact in which the most complex model appears best because it is over-fitted. We therefore suggest that a reasonable strategy for selecting variables to include in SDMs is to begin with geographic coordinates (and a sampling effort covariate such as checklist length, if appropriate), and then add environmental variables, limiting the number of environmental variables based on sample size to avoid overfitted models. Many studies have noted the benefits of explicitly including space in SDMs (Lennon, 2000; Dormann, 2007; Beale et al., 2010); we suggest taking advantage of the nearly universal presence of spatial autocorrelation in species distributions by including geographic coordinates as part of a “base model” to which environmental covariates can be added. Our goal in including geographic coordinates as predictor variables was to take advantage of spatial structure for prediction (Bahn & McGill, 2007), rather than to better estimate the effects of predictor variables or control for pseudo-replication or spatial structure in the errors (Beale et al., 2010). Using geographic coordinates as predictor variables has the advantage that decent predictive models can likely be constructed even for species for which the most relevant environmental drivers of distribution are not known.

The good performance of the spatial models is encouraging for predictive models, but discouraging for attempts to identify biologically meaningful environmental drivers of distributions. Geographic coordinates were regularly in the top half of variables ranked by importance for our SDMs (Figs. S3-S4), highlighting the fact that the usefulness of a variable for prediction does not provide insight about whether the variable is a causative determinant of distribution.

Dispersal is likely an important determinant of distributions for some millipede species. Millipedes may be dispersed long distances by humans in soil and plant material, but once established in new locations, local dispersal may be much slower (Baker, 1978), though some species can be mobile and disperse readily (David & Handa, 2010). Limited local dispersal could lead to patchy distributions in which large parts of suitable environmental space are not occupied, obscuring any signal of environmental suitability from SDMs. Models using geographic coordinates as predictors may be better able to capture such patchy distributions, and may predict well when interpolating between sampling locations, even though they can not extrapolate to new geographic areas.

The ranking of the predictor variables by relative importance in the random forest models changed when models were trained with spatially under-sampled rather than raw data (Figs. S3-S4). Variable importance measures (and partial dependence plots) from our SDMs can be used to suggest hypotheses about factors that influence the distribution and/or detectability of species (Kelling et al., 2009). Knowledge of the life history and ecology of millipedes in Ireland is patchy, with almost nothing known about some species (Lee, 2006). Hypotheses generated by exploring our variable importance and partial dependence plots can provide a basis for future confirmatory analyses.

However, because the variable importance rankings for most species changed when using spatially under-sampled data, we have low confidence in interpreting variable importance rankings in terms of ecology The raw data were opportunistically collected, and the spatially sub-sampled data were therefore also opportunistic (since they were a subset of the original data). The goal of spatial under-sampling was to improve prediction performance rather than to improve insight about ecological or physiological determinants of occurrence; we do not know of any *a priori* reason to think that either the raw or under-sampled data produced more ecologically correct inferences about the ecological importance of predictor variables. For some variables, including checklist length (discussed below), variable importance rankings were similar for models trained with both datasets, providing somewhat stronger evidence for the relative importance of those variables in predicting our data.

We used checklist length as a proxy for sampling effort, but checklist length probably also varied with factors not related to sampling effort, including species richness (Warton et al., 2013). Checklist length can be used as a sampling effort covariate in the detection sub-model within hierarchical occupancy-detection models (MacKenzie et al., 2002), or to account for detectability in non-hierarchical models (Szabo et al., 2010; Isaac et al. 2014). The intuition we followed is the same. To the extent that checklist length successfully captured variability in sampling effort, the predictions generated using a standardized checklist length (Fig. 6a-b, Figs. S11-S15) represent the relative probability of recording the focal species when sampling effort is constant in each grid cell. Checklist length was the most important variable in models for three out of our six focal species (Figs. S3-S4). Partial dependence plots showed that relationships between checklist length and relative probability of the focal species being recorded were generally positive, with probability of the species being recorded increasing more quickly when checklist length was small, as expected (Fig. 5f, Figs. S5-S10).

**Figure 6:**
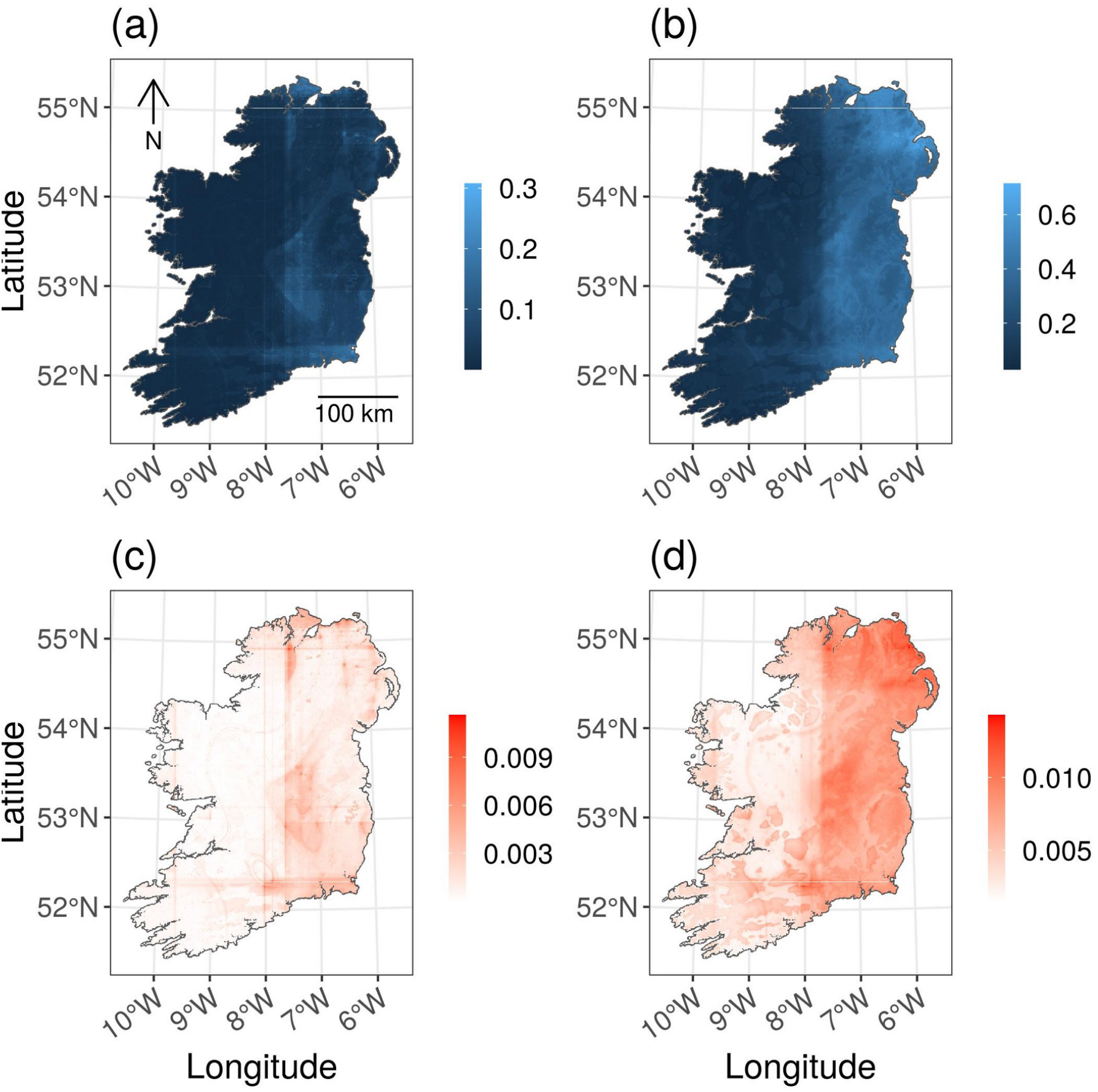
Predicted distribution of the millipede *Ommatoiulus sabulosus* in Ireland. The top maps show mean predicted relative probability of detecting *O. sabulosus* on a checklist of length two, from the *ENVIRONMENT + COORDINATES + SEASON + LIST LENGTH* model trained with (a) raw data and (b) spatially under-sampled data. Bottom maps show: (c) standard error of the mean predictions from (a); and (d) standard error of the mean predictions from (b). The standard errors of the predictions in each grid cell (c) and (d) show how much model predictions varied based on which records were included or excluded from the cross-validation training dataset. Structure from the predictor variables is visible in model predictions and standard errors, for example as vertical and horizontal lines (artifacts of the geographic coordinate predictor variables) and contour lines (e.g. in the southeast quadrant of (a), an artifact of the precipitation variable).

Checklist length is not a perfect proxy for sampling effort, because it confounds sampling effort and the number of species available to be recorded (Warton et al., 2013). The number of species available for detection is probably not constant through the year in many locations in Ireland because of seasonal patterns in detectability. Likewise, the total number of millipede species present in 1 km x 1 km grid cells is almost certainly not constant across all grid squares in Ireland. Checklist length is therefore determined by sampling effort, species richness (which varies spatially), and detectability (which varies seasonally). In our data, checklist length was not correlated with any of the environmental predictor variables (absolute value of Spearman’s correlation coefficient was always < 0.1 for correlations between checklist length and other predictor variables), and there were no obvious geographic patterns in checklist length (Fig. S16). Most locations in Ireland probably have multiple millipede species available for detection at most times of year (i.e. adults of multiple species present), but, as is common in biological records datasets, many of the checklists in our dataset had checklist lengths of one (37% of checklists) or two (23% of checklists). We suspect that checklists of only one or two species resulted from limited sampling effort rather than intensive surveys in locations with only one or two species. Our use of checklist length provided no information about cases in which surveys were conducted but no species were found. There may be times of the year and/or 1 km^2^ grid squares in which there truly are no millipedes to be found (e.g. in grid squares dominated by bog), but our training data do not contain information about those areas.

## 5 Conclusion

Species distribution models for rare millipede species had better prediction performance according to most metrics when the training data had been spatially under-sampled by discarding non-detection data to improve class balance and spatial bias. Models trained with spatially under-sampled data had worse specificity (true negative rate) than models trained with raw data, but the decrease in specificity was small for the rarest species and was accompanied by large improvements in sensitivity (true positive rate).

A comparison of models using geographic coordinates as predictor variables and models using environmental predictor variables showed that neither set of variables always outperformed the other. Notably, the distribution of *O. sabulosus* was better predicted using geographic coordinates rather than environmental variables. Models combining both geographic coordinates and environmental variables as predictors were consistently among the best performing models.

We tested spatial under-sampling using a smaller, sparser dataset than was used in previous tests (Robinson et al., 2018; Robinson et al., 2020). Modeling distributions using sparse datasets is beneficial when it is difficult or expensive to collect additional observational data about species occurrence. Traditional survey methods (including citizen science) will likely never produce large occurrence datasets for taxa that are small, difficult to identify, and/or non-charismatic (though other approaches including eDNA may be able to provide large amounts of data about such taxa). Our results suggested that spatial under-sampling can be used to improve SDMs of rare, non-charismatic, poorly sampled taxa, including invertebrates, for which biological recording effort is limited.

## Acknowledgments

We thank Tomás Murray and the Irish National Biodiversity Data Centre (NBDC) for advice about the biological records data, and Declan Doogue for information about millipedes in Ireland. We thank the British Myriapod and Isopod Group and the many citizen scientists who collected and contributed data.

## Notes

***Funding Statement*** This publication emanated from research supported in part by Science Foundation Ireland (grant number 15/IA/2881). The opinions, findings and conclusions or recommendations expressed in this material are those of the author(s) and do not necessarily reflect the views of the Science Foundation Ireland. This research used CORINE data made available with funding by the European Union.

https://doi.org/10.15468/dl.833k97

https://github.com/wgaul/maps_of_ignorance/tree/master/millipede_maps_of_ignorance

### Competing Interest Statement

The authors have declared no competing interest.

