## Supplemental Figures for "Modelling the distribution of rare invertebrates by correcting class imbalance and spatial bias"

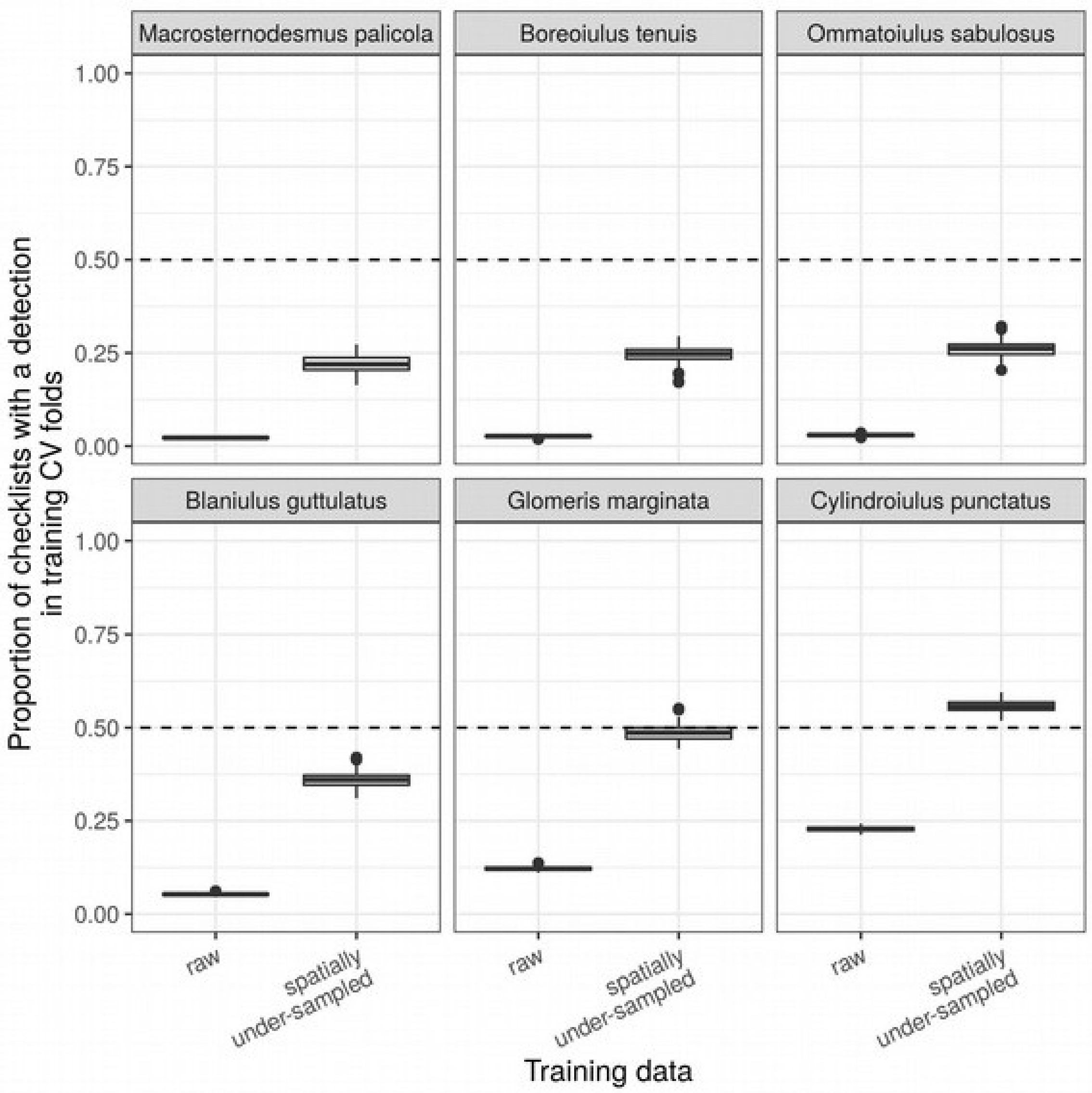


**Figure S1:** Class balance in raw (left) and spatially under-sampled (right) data used to train species distribution models for six millipede species in Ireland. Perfect class balance would be represented by a value of 0.5 for the proportion of checklists with a detection (horizontal dashed line). Each boxplot shows the proportion of checklists with a detection for that species in 99 randomly re-sampled training datasets. Spatial under-sampling improved the class balance for all species. For the most common species in the dataset, *Cylindroiulus punctatus*, spatial under-sampling resulted in more detections than non-detections in the training data, but the proportion was closer to 0.5 than the proportion in the raw data, meaning class balance was improved.

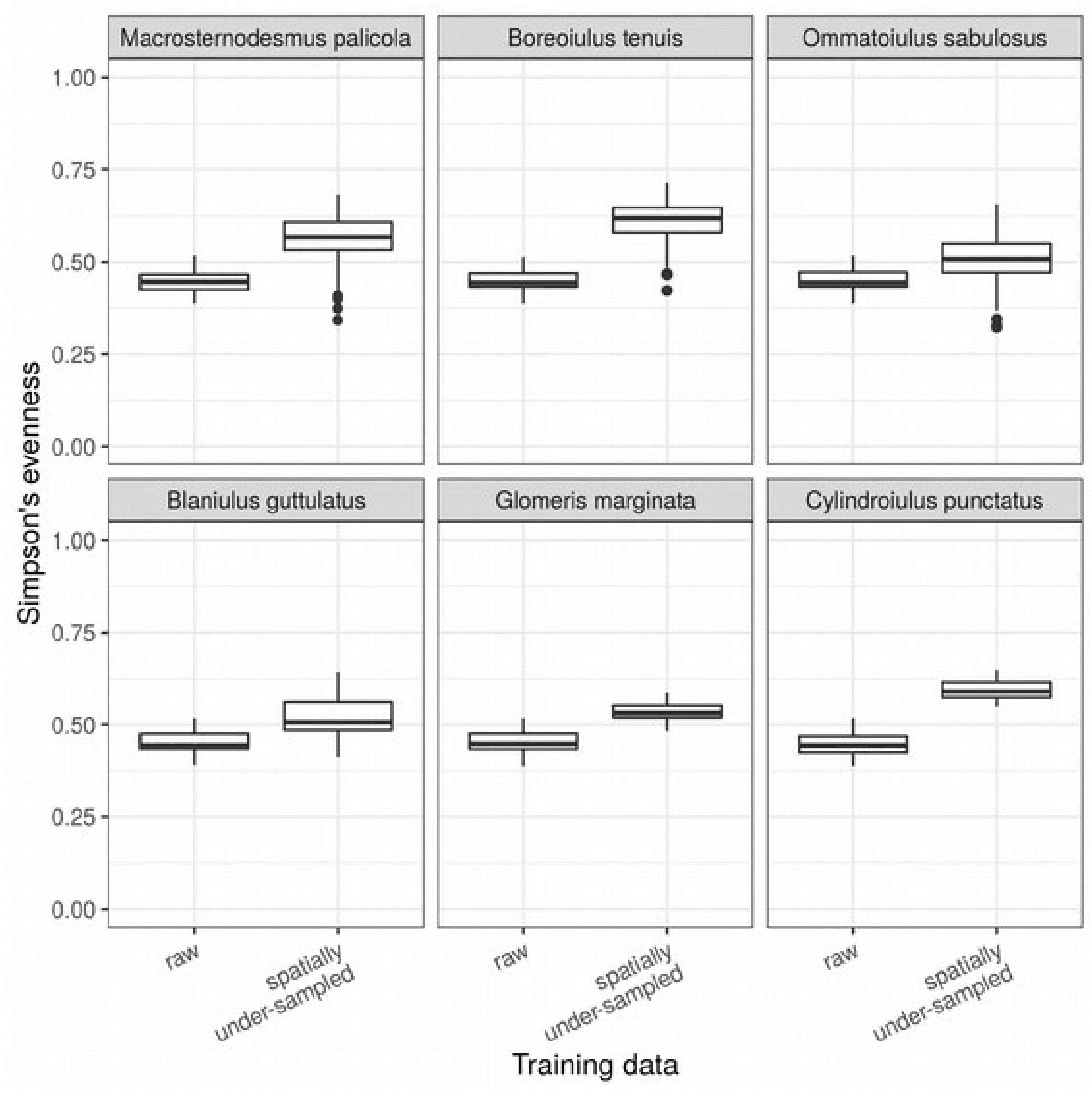


**Figure S2:** Spatial evenness of raw and spatially under-sampled training data used to fit species distribution models for six millipede species in Ireland. Spatial evenness was measured using Simpson's evenness to quantify the evenness in the number of checklists in 30 x 30 km grid squares covering Ireland. Spatial under-sampling improved the spatial evenness of the training data at this spatial scale. Each boxplot shows the distribution of Simpson's evenness values from 132 different random placements of the 30 x 30 km grid.

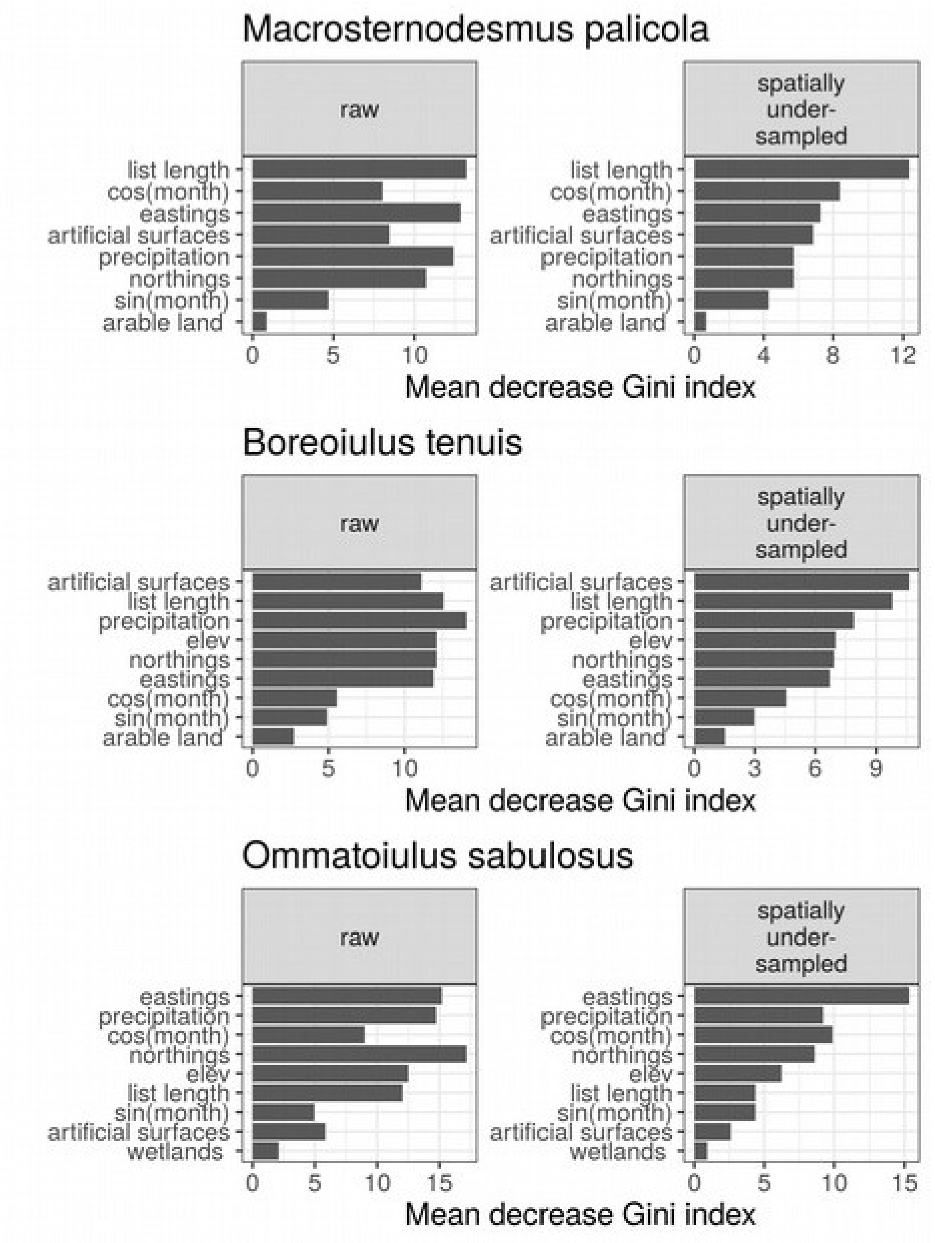


**Figure S3:** Variable importance from random forest species distribution models of millipede species in Ireland, trained with raw and spatially under-sampled data. Variable importance was measured as the decrease in node impurity (measured using the Gini index) from splitting on each variable. Variable importance was averaged over all trees within models and over all 99 model runs for each species/training data combination. Spatially under-sampling the training data resulted in different rankings of variable importance for some species, including differences in which variable was ranked most important. The ranking of the least important variables generally did not change.

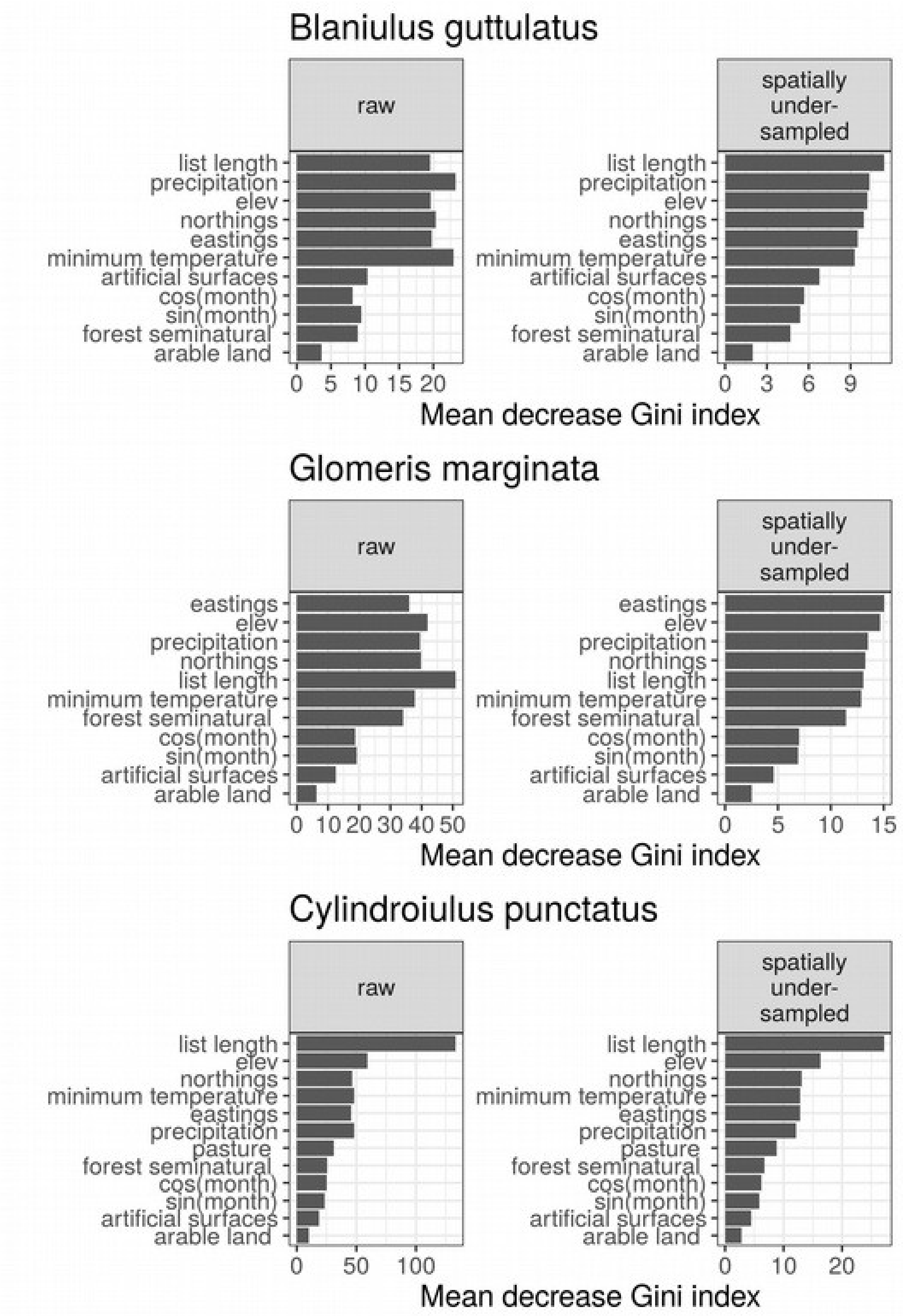


**Figure S4:** Variable importance from random forest species distribution models of millipede species in Ireland. Details as for Fig. S3.

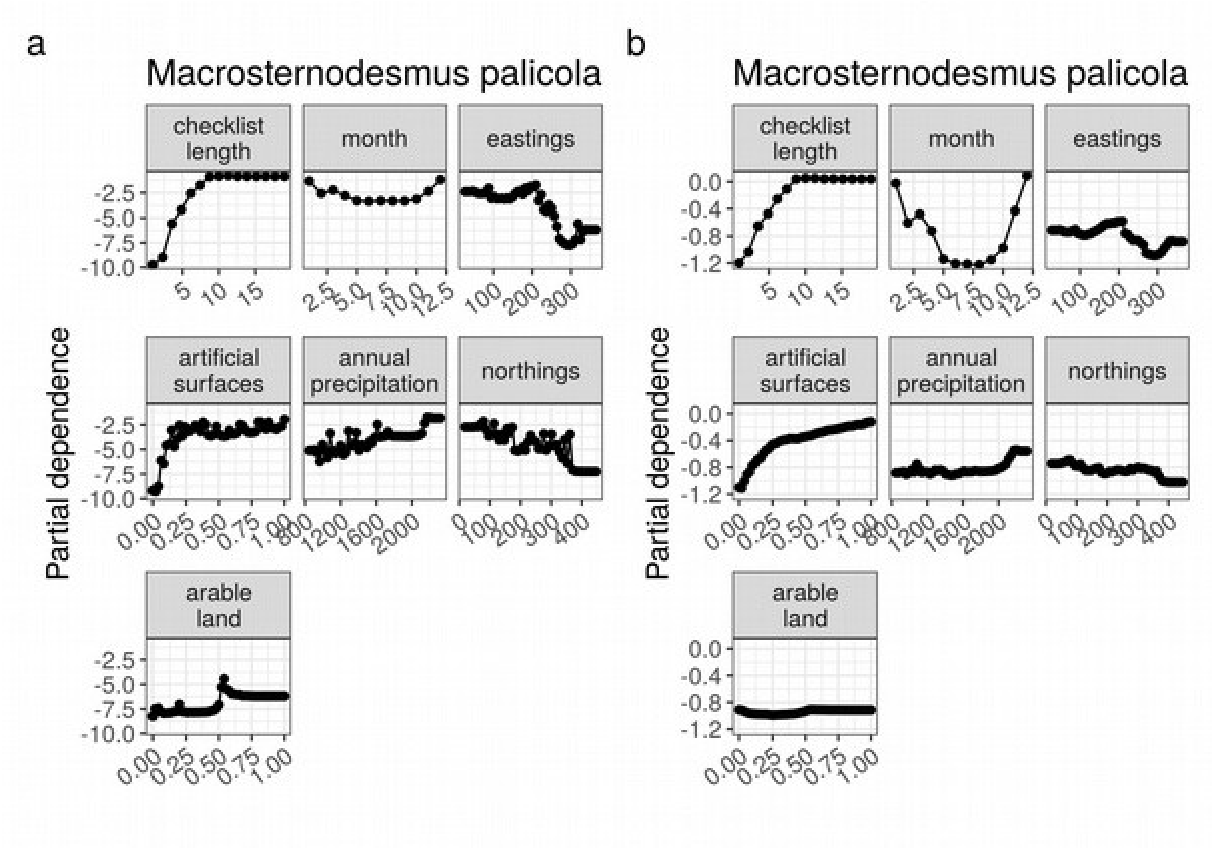


**Figure S5:** Partial dependence plots from random forest species distribution models showing the effect of each predictor variable on the probability of detecting *Macrosternodesmus palicola* on a checklist. Results are shown from the most complex model (ENVIRONMENT + COORDINATES + SEASON + LIST LENGTH), trained with raw (a) and spatially under-sampled (b) data. Variables are arranged from highest to lowest importance based on variable importance measures from models trained with spatially under-sampled data.

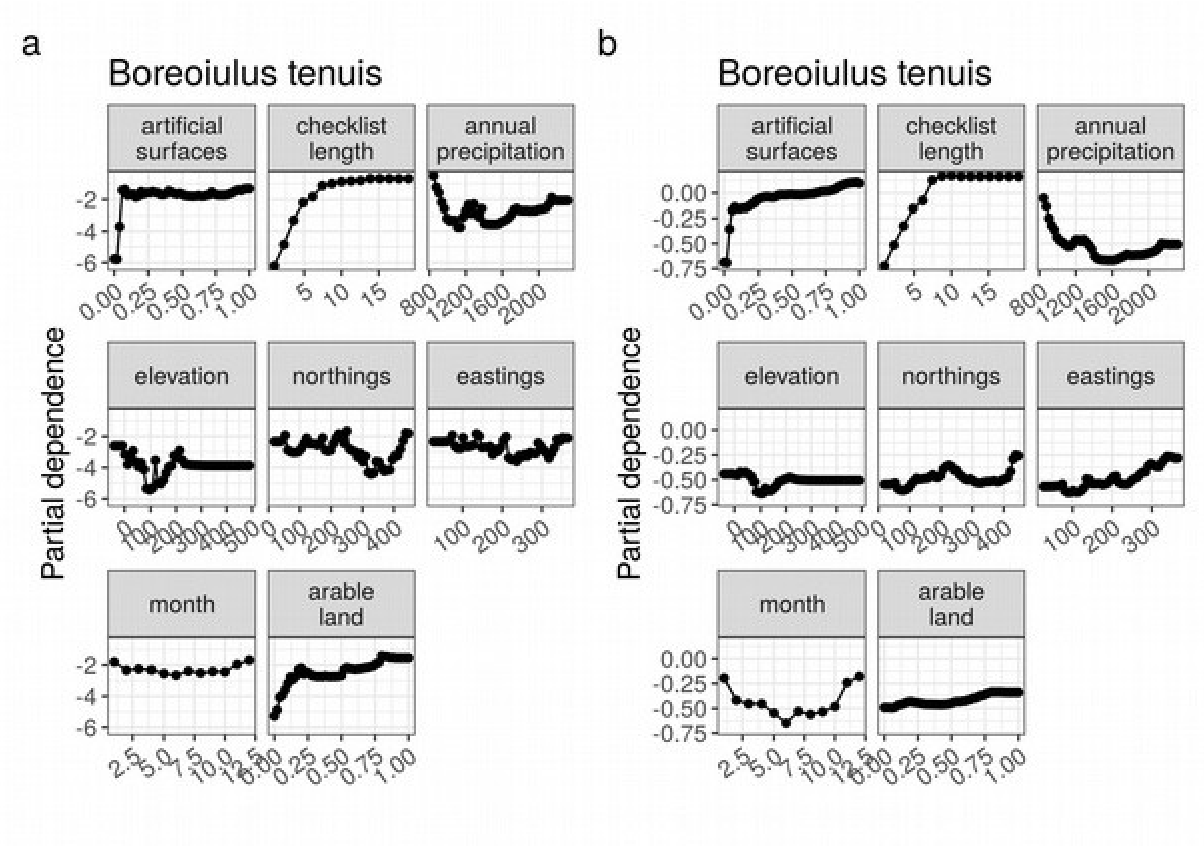


**Figure S6:** Partial dependence plots from random forest species distribution models showing the effect of each predictor variable on the probability of detecting *Boreoiulus tenuis* on a checklist.  Details are as for Fig. S5.

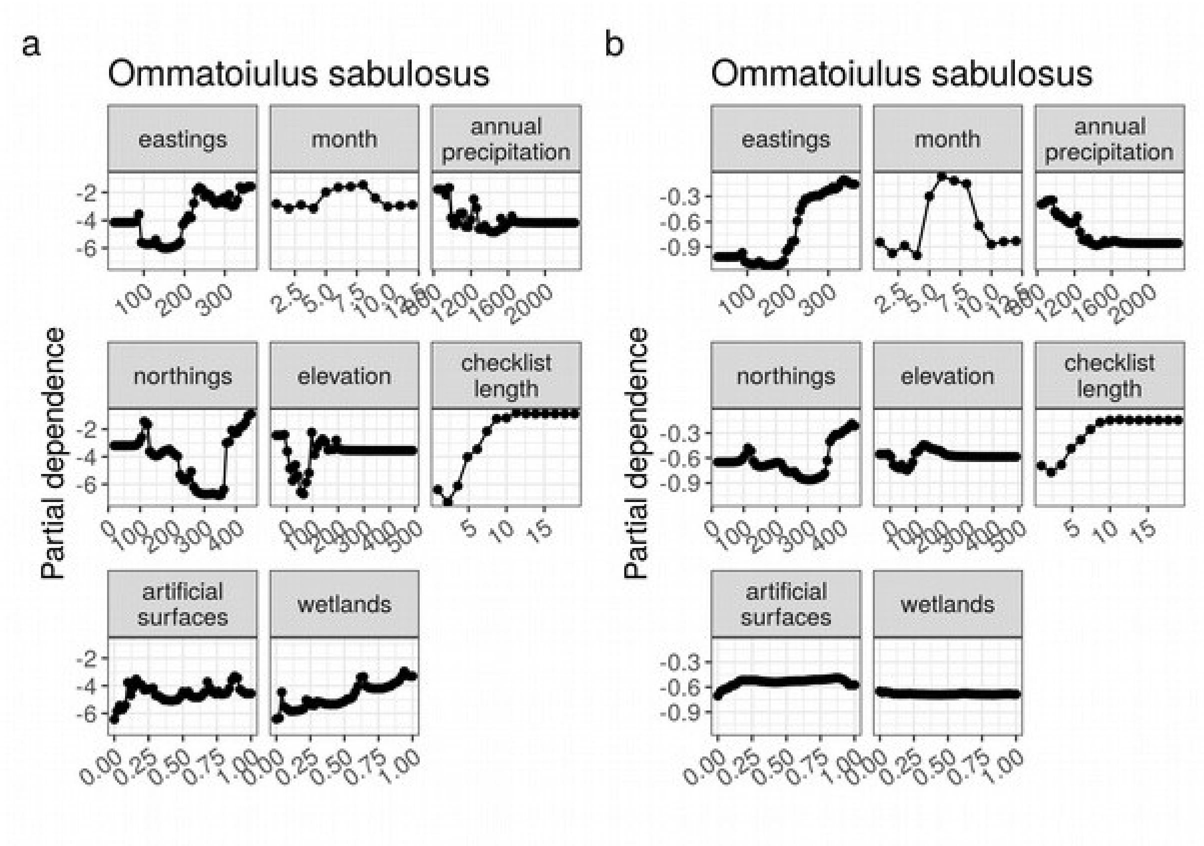


**Figure S7:** Partial dependence plots from random forest species distribution models showing the effect of each predictor variable on the probability of detecting *Ommatoiulus sabulosus* on a checklist. Details are as for Fig. S5.

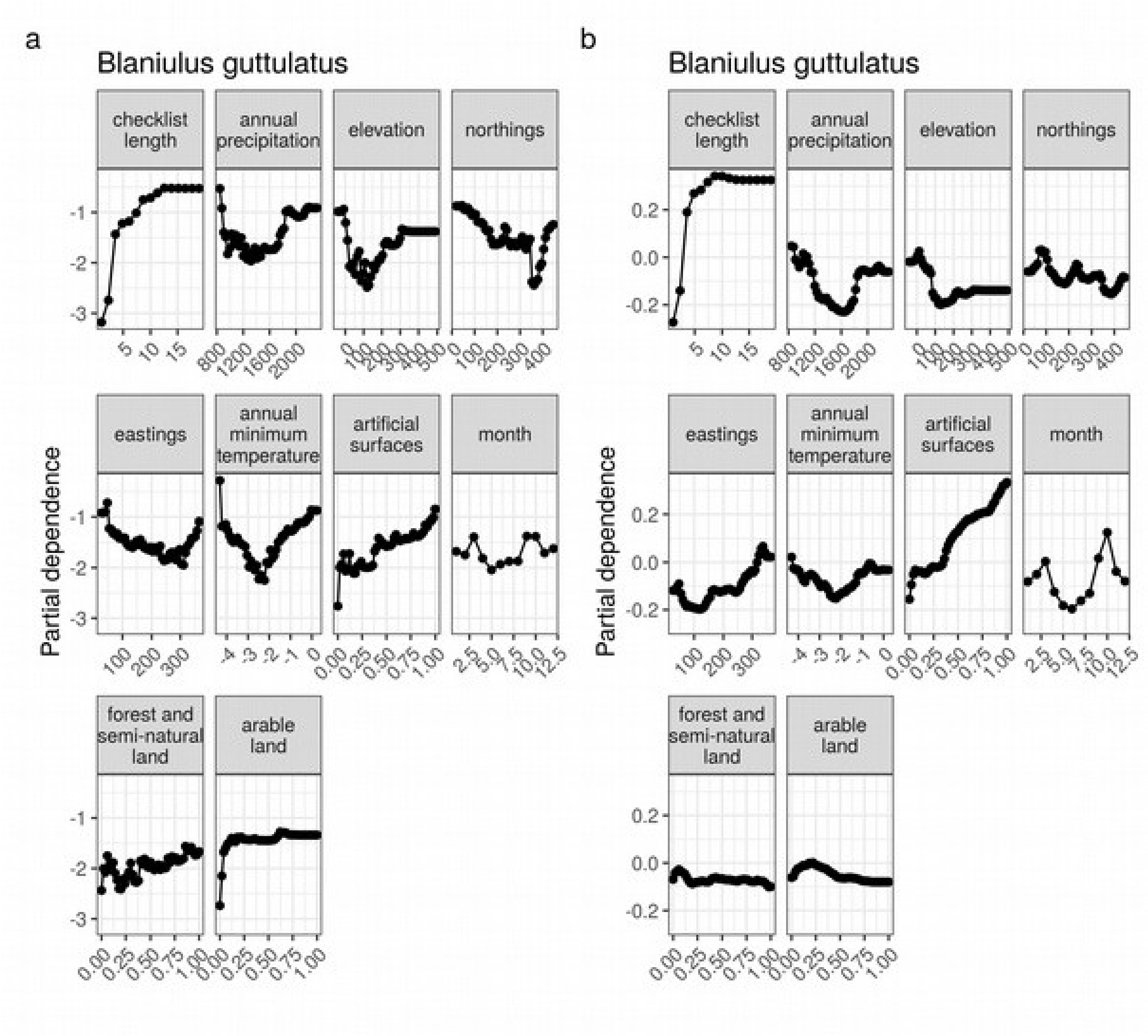


**Figure S8:** Partial dependence plots from random forest species distribution models showing the effect of each predictor variable on the probability of detecting *Blaniulus guttulatus* on a checklist. Details are as for Fig. S5.

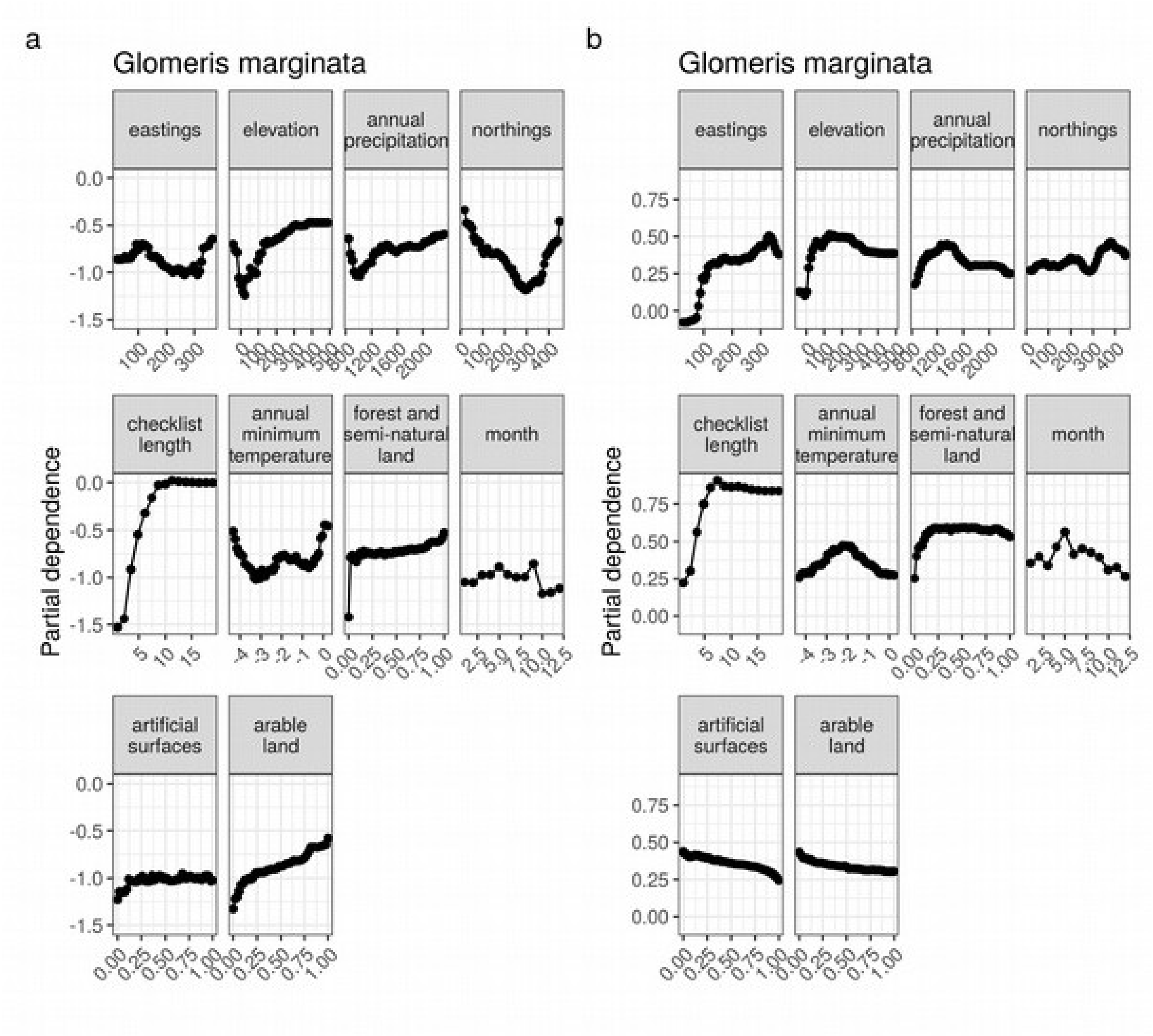


**Figure S9:** Partial dependence plots from random forest species distribution models showing the effect of each predictor variable on the probability of detecting *Glomeris marginata* on a checklist. Details are as for Fig. S5.

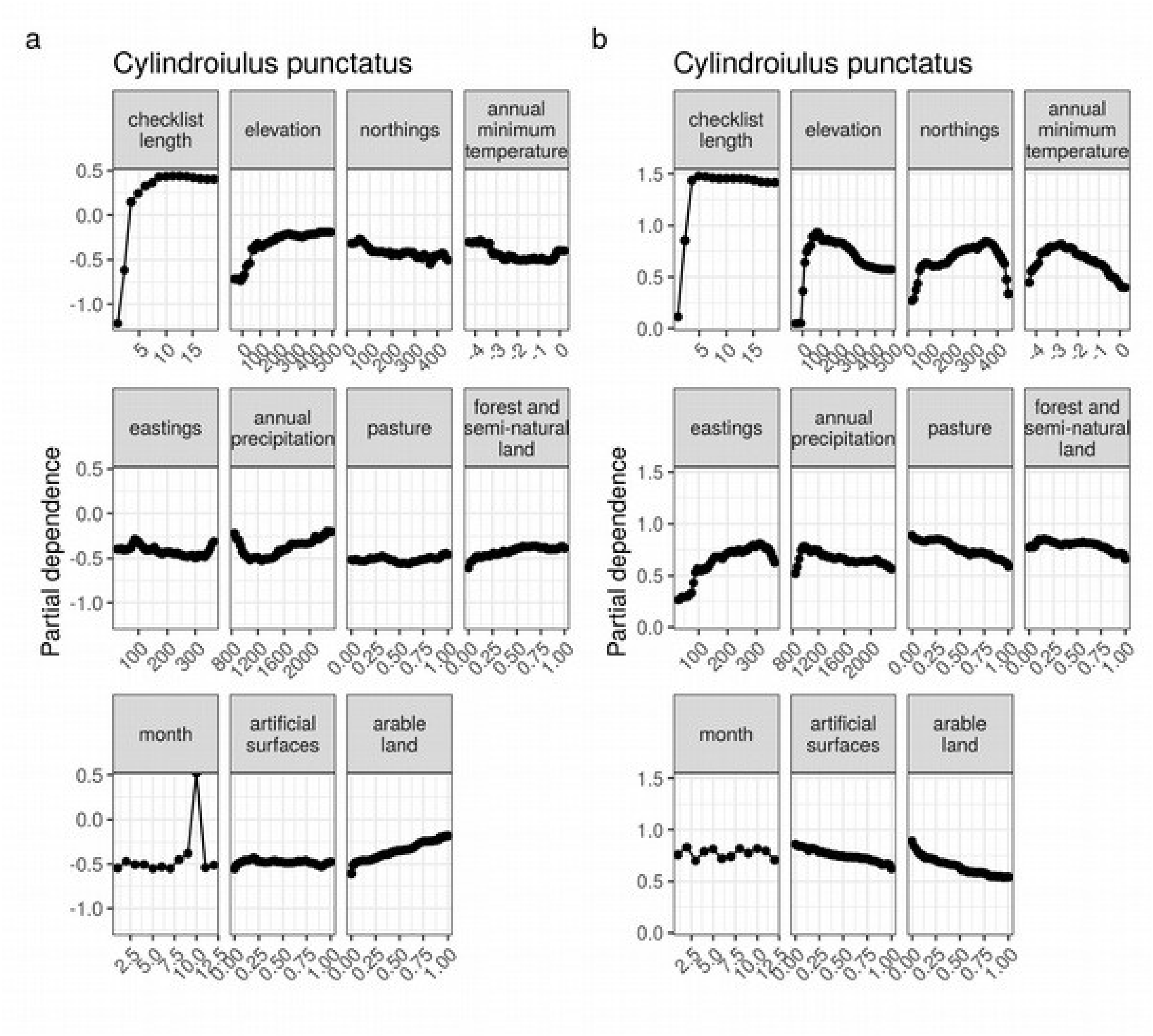


**Figure S10:** Partial dependence plots from random forest species distribution models showing the effect of each predictor variable on the probability of detecting *Cylindroiulus punctatus* on a checklist. Details are as for Fig. S5.

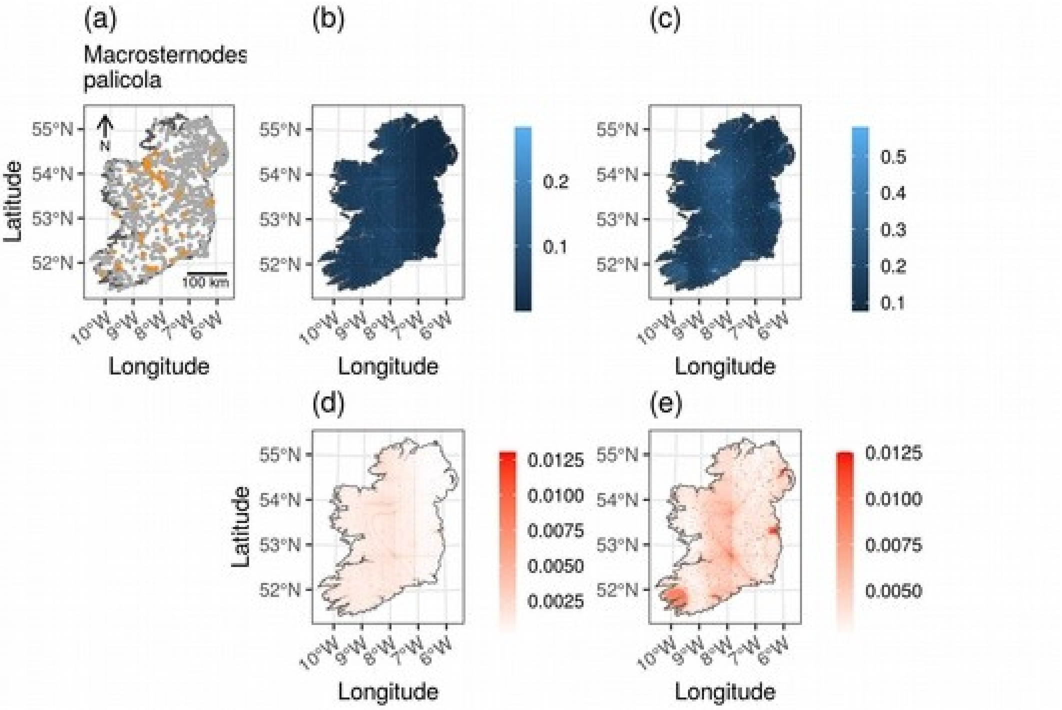


**Figure S11:** The distribution of the millipede *Macrosternodesmus palicola* in Ireland. Maps show: (a) detections (orange points) and non-detections (grey points) of the species on checklists; (b) mean predicted relative probability of detecting *M. palicola* on a checklist of length two, from the *ENVIRONMENT* + *COORDINATES* + *SEASON* + *LIST LENGTH* SDM trained with raw data; (c) the same as (b) but when model was trained with spatially under-sampled data; (d) standard error of the mean predictions from (b); and (e) standard error of the mean predictions from (c). The predicted distributions (b) and (c) show mean predictions in 1 $km^2$ grid squares from 99 replicates of the model. The standard errors of the predictions in each grid cell (d-e) show how much model predictions varied based on which records were included or excluded from the cross-validation training dataset. Predicted probabilities are relative rankings of probability of detection, but are not calibrated to true probabilities of occurrence. It is meaningful to compare relative probabilities within a model (e.g. to determine whether the focal species is more likely to occur in one grid square than another), but it is not meaningful to compare relative probabilities between models or between species (i.e. it is not possible to determine whether one species is more likely than another species to occur in a particular grid square).

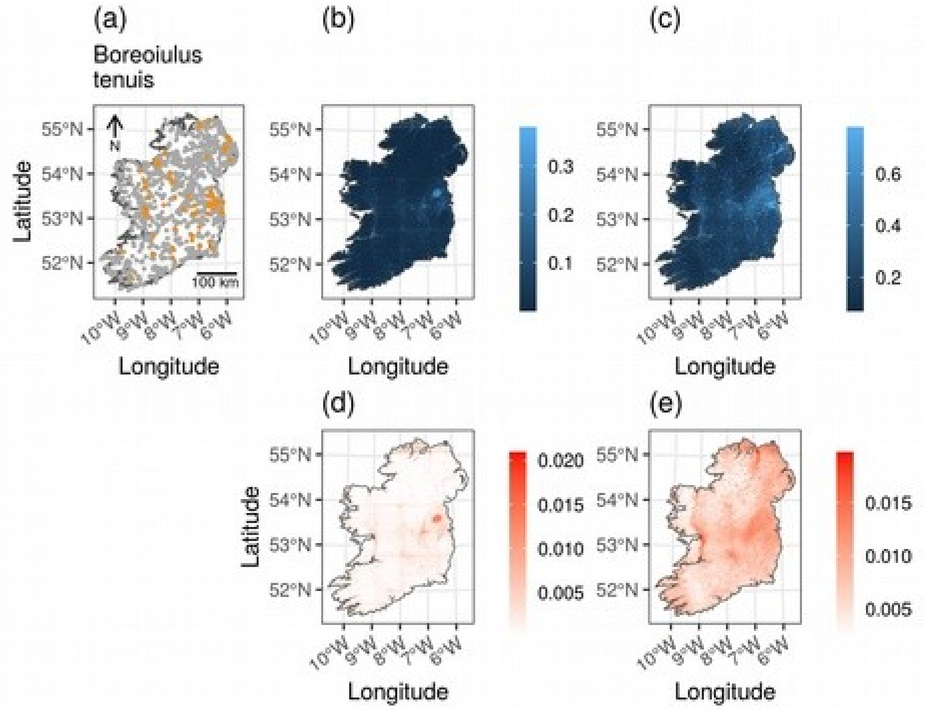


**Figure S12:** The distribution of the millipede *Boreoiulus tenuis* in Ireland. Details are as for Fig. S11.

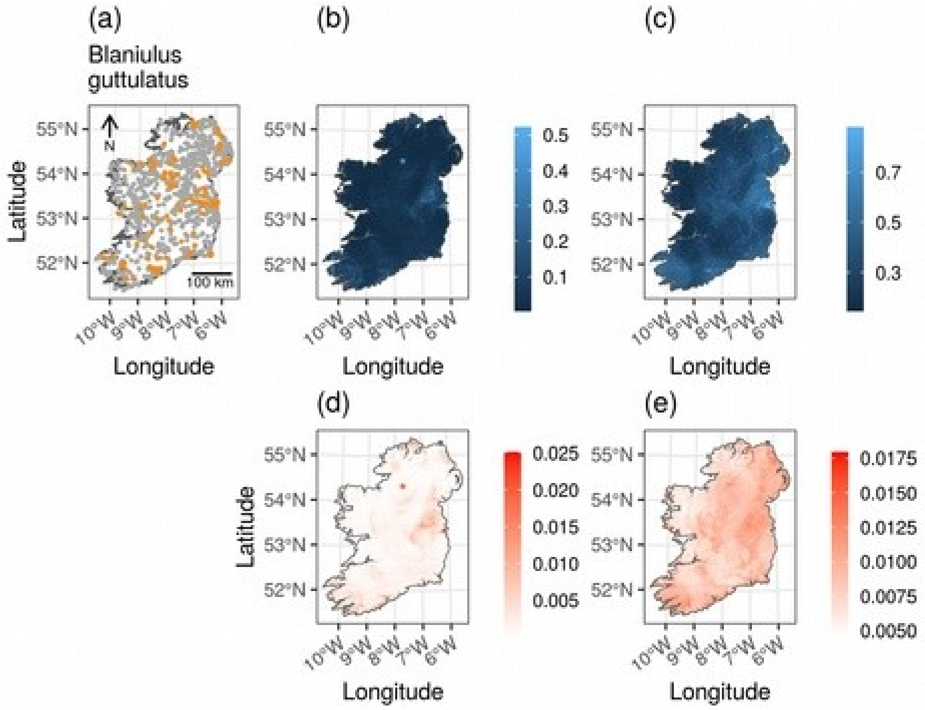


**Figure S13:** The distribution of the millipede *Blaniulus guttulatus* in Ireland. Details are as for Fig. S11.

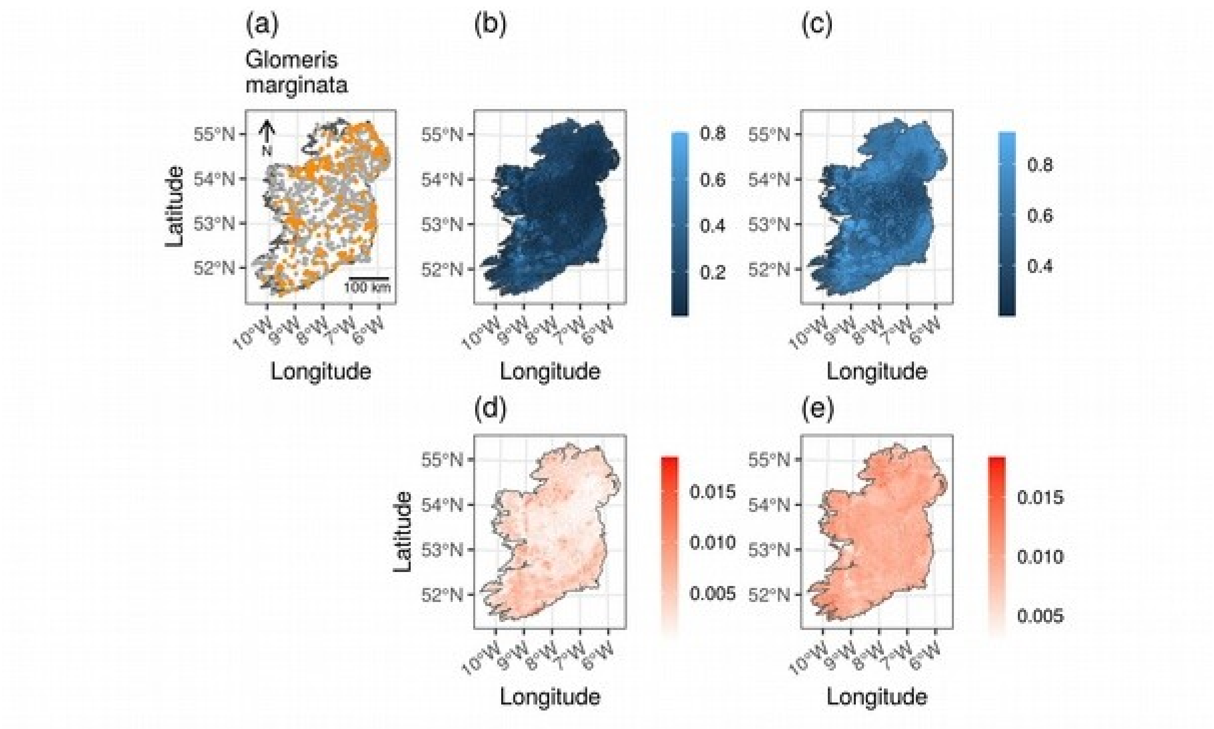


**Figure S14:** The distribution of the millipede *Glomeris marginata* in Ireland. Details are as for Fig. S11.

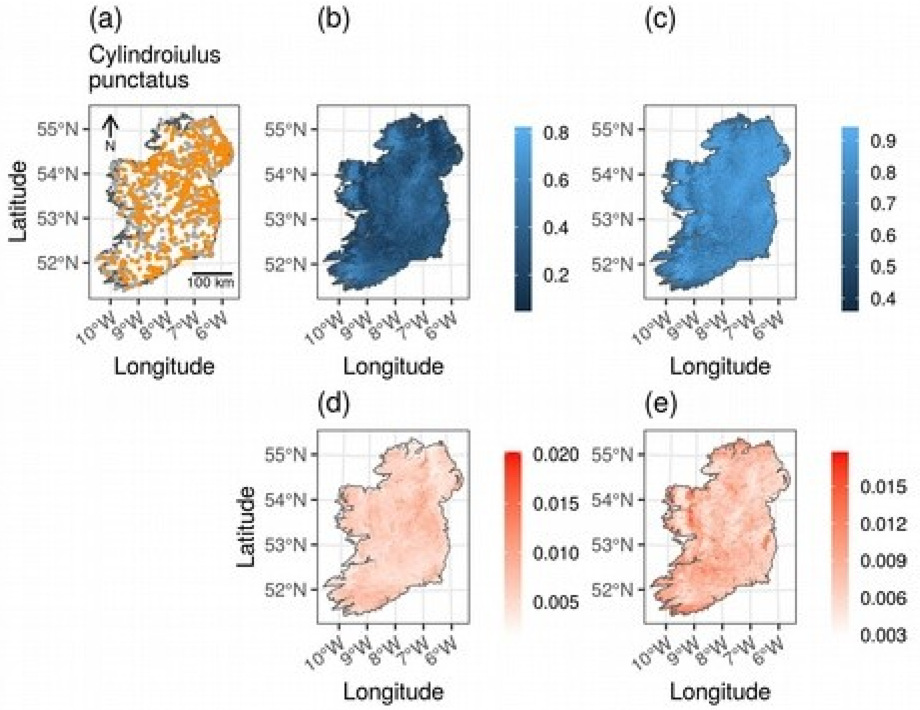


**Figure S15:** The distribution of the millipede *Cylindroiulus punctatus* in Ireland. Details are as for Fig. S11.

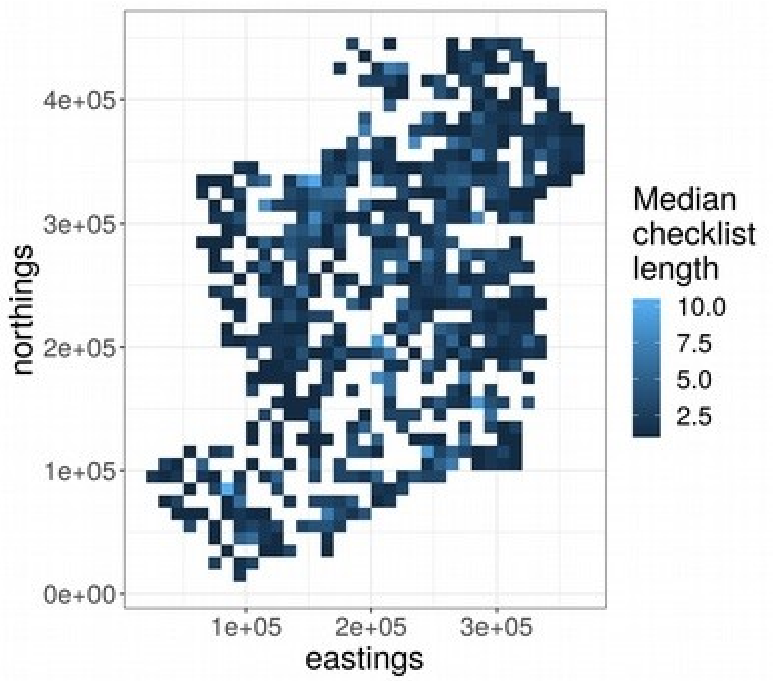


**Figure S16:** Median checklist length for millipede checklists in 10 x 10 km grid squares in Ireland. Only grid squares with at least one checklist are shown. Coordinates are eastings and northings of the TM75 Irish Grid Reference system.
