## Appendix S1 for "Modelling the distribution of rare invertebrates by correcting class imbalance and spatial bias"

### Appendix S1 – Supplementary methods

#### Month transformation

To allow for the cyclical nature of variation in detectability with month, we used variations of cosine and sine transformations of month (James, 2011) to create two separate transformed month variables that we provided to the random forest SDMs. The transformed month variables allowed December and January to be close in data space, which allowed the models to capture annual periodic changes in detection probability (Fig. 5). The transformed variables used in models were:

$$Msin_i = \sin\left(\frac{2\pi M_i}{12}\right)$$

and

$$Mcos_i = \cos\left(\frac{2\pi M_i}{12}\right)$$

where $M_i$ is the month ($i$ indexes the month with values of 1 through 12).

#### Partial dependence measures

Partial dependence for predictor variables other than month was calculated using the 'partialPlots' function in the 'randomForest' R package (Lias & Wiener 2002). For month, the variables included in the models were $M_{sin}$ and $M_{cos}$, not raw month, but the partial dependence relationship that is most biologically interpretable is the dependence of species detection on month. Therefore, we calculated the partial dependence on month by calculating the partial dependence on the predictor variables $M_{sin}$ and $M_{cos}$ representing actual months. Following Hastie, Tibshirani, and Friedman (2013, section 10.13.2), we calculated partial dependence on the predictor variables $M_{sin}$ and $M_{cos}$ for months $m$ as

$$f(m) = \frac{1}{n}\sum_{i=1}^{n}\log\left(p\left(Msin_m, Mcos_m, x_{Ci}\right)\right) - \frac{\frac{1}{n}\sum_{i=1}^{n}\log\left(p\left(Msin_m, Mcos_m, x_{Ci}\right)\right) + \frac{1}{n}\sum_{i=1}^{n}\log\left(1 - p\left(Msin_m, Mcos_m, x_{Ci}\right)\right)}{2}$$

where $p$ is the predicted probability of recording the focal species on a checklist with predictor variables $M_{sin}$, $M_{cos}$, and $x_{Ci}$. n is the number of training data checklists, $X_C$ is the complement set of predictor variables other than $M_{sin}$ and $M_{cos}$, and $x_{Ci}$ are the values of the predictor variables in $X_C$ from the $i_{th}$ checklist in the training data.
